# Mapping the anatomical and transcriptional landscape of early human fetal ovary development

**DOI:** 10.1101/2023.09.26.558771

**Authors:** Sinead M McGlacken-Byrne, Ignacio del Valle, Theodoros Xenakis, Ian C Simcock, Jenifer P Suntharalingham, Federica Buonocore, Berta Crespo, Nadjeda Moreno, Danielle Liptrot, Paola Niola, Tony Brooks, Gerard S Conway, Mehul T Dattani, Owen J Arthurs, Nita Solanky, John C Achermann

**Affiliations:** Genetics and Genomic Medicine Research and Teaching Department, UCL Great Ormond Street Institute of Child Health, University College London, London WC1N 1EH, United Kingdom; Department of Clinical Radiology, Great Ormond Street Hospital for Children NHS Foundation Trust, London, WC1N 3JH, United Kingdom; Population, Policy and Practice Research and Teaching Department, UCL Great Ormond Street Institute of Child Health, University College London, London, WC1N 1EH, United Kingdom; NIHR Great Ormond Street Biomedical Research Centre, London, WC1N 1EH, United Kingdom; Developmental Biology and Cancer Research and Teaching Department, UCL Great Ormond Street Institute of Child Health, University College London, London WC1N 1EH, United Kingdom; UCL Genomics, Zayed Centre for Research, UCL Great Ormond Street Institute of Child Health, University College London, London WC1N 1DZ, United Kingdom; Institute for Women’s Health, University College London, London, WC1E 6AU, United Kingdom

## Abstract

The complex genetic mechanisms underlying human ovary development can give rise to clinical phenotypes if disrupted, such as Primary Ovarian Insufficiency and Differences of Sex Development. Through a clinically-focused lens, we combine single-nuclei RNA sequencing, bulk RNA sequencing, and micro-focus computed tomography to elucidate the anatomy and transcriptional landscape of the human fetal ovary across key developmental timepoints (Carnegie Stage 22 until 20 weeks post conception). We show the marked growth and distinct morphological changes within the fetal ovary at the critical timepoint of germ cell expansion, and demonstrate that the fetal ovary becomes more transcriptomically distinct from the testis with age. We describe novel ovary developmental pathways, relating to neuroendocrine signalling, energy homeostasis, mitochondrial networks, piRNA processes, and inflammasome regulation. We define transcriptional regulators and candidate genes for meiosis within the developing ovary. Together, this work advances our fundamental understanding of human ovary development and clinical ovarian insufficiency phenotypes.

## Introduction

In humans, testes or ovaries develop from the bipotential gonad at around 42 days post conception (dpc)^1^. It is well established that testis determination is driven by expression of the male-typical testis-determining factor, *SRY*, at this timepoint, leading to upregulation of *SOX9* and a cascade of downstream pro-testis pathways^2^. In contrast to this, an “ovary-determining factor” similar to SRY has not been identified, and for many years ovary development was considered a largely “passive” process occurring in the absence of SRY^3^. A key differentiating feature between ovary and testis development are the biological processes of germ cell expansion and meiotic entry that occur in the fetal ovary. These primordial follicles – representing an individual’s entire ovarian reserve and the germ cell pool for the next generation – are suspended in meiosis I until puberty many years later. The advent of high throughput transcriptomic approaches has begun to characterize ovary development and its key biological processes in previously unparalleled detail, revealing the process to be multigenic, complex and, still, only partially understood.

Initial microarray studies of gene transcription in developing mouse gonads were first reported at scale in 2007 and introduced the concept that a similar number of genes were differentially expressed in the ovary compared to the testis^4,5^. Many of these genes were proposed to antagonize testis development or were involved in meiosis. Analysis of developing human gonad tissue eventually followed. An array-based RNA-sequencing atlas confirmed that the human fetal ovary had a discrete transcriptomic signature with clearly upregulated ovary-enriched genes^6^. Some of these had known roles in the developing ovary, such as *NANOG* and *POU5F1* (OCT3/4), whereas the roles of others remained elusive. Pathway enrichment analysis revealed little further insight with most genes not annotated to known biological functions.

More recently, a bulk RNA-sequencing (bulk RNA-seq) study of human developing gonadal tissue between 6 and 17 weeks post conception (wpc) demonstrated several expected features, such as marked upregulation of meiotic transcripts, but also some novel findings, such as an enrichment of non-coding RNAs and genes with no clear link to ovary development (e.g., neuropeptide *TAC1* and neurexin *NRXN3*)^7^. A limited number of single-cell RNA-sequencing (scRNA-seq) studies have recently built upon these foundations to demonstrate important cell-cell interactions, ovary developmental pathways, and gonadal somatic and germ cells crosstalk (such as FGF9-FGFR2 and PROS1-TYRO3)^8,9^. New technologies have facilitated further multi-modal -omic analyses; particularly, the integration of single cell ATAC-sequencing (assay for transposase-accessible chromatin with sequencing) with single-cell data has elucidated novel post-transcriptional regulatory layers in the developing ovary^8,9^. Lastly, larger-scale analyses involving multiple species demonstrate *human-specific* aspects of gonadal development; for example, the observation that expression of *KLF4, ELF3,* and *ZNF581* is highly specific to the human germ cell specification programme^9^. These findings highlight the potential limitations of applying animal RNA-seq data to humans and the importance of studying human gonadal tissue where possible.

Using bulk and single-cell technologies in isolation to assess transcriptomic expression in tissues has limitations. Single nuclei RNA-seq (snRNA-seq) provides granular transcriptomic detail, localizes gene expression to individual cell populations, and reconstructs cellular differentiation networks *in silico*. However, sensitivity can be low and only a proportion of the transcriptome may be captured. Bulk RNA-seq captures a higher proportion of transcriptomic activity, but sensitivity is also compromised in that low-level expression from small but important cell populations can be lost^10^. Additionally, while large-scale RNA sequencing atlases provide vital information about transcriptional landscapes, the potential clinical impact of these findings is not always explored. If the genetic mechanisms that span a female reproductive lifecourse from fetal life to adulthood are disrupted, ovarian dysfunction can result. This manifests in a range of clinical phenotypes including ovarian insufficiency occurring in isolation (Primary Ovarian Insufficiency, POI), as a feature of an extra-ovarian syndrome (e.g., Perrault Syndrome), or as part of a Difference of Sex Development (DSD) (e.g., aromatase [*CYP19A1*] deficiency). POI affects up to 1% of women and results in infertility, a need for lifelong hormone replacement therapy, and medical co-morbidities including cardiovascular and bone health consequences. DSD is less common and represents several conditions diagnosed at different timepoints across the reproductive lifecourse. Next generation sequencing approaches have increased the genetic diagnostic yield in reproductive disorders – for example, a genetic cause is identified in over 20% of women with POI and 30% of those with DSD, although most remain unexplained^11,12^. A granular understanding of the pathogenic mechanisms responsible for the genetically heterogenous condition of ovarian insufficiency is key for an individualized approach to clinical management and reproductive counseling. Further, this knowledge may be harnessed to develop novel fertility preservation techniques, particularly if the pre-symptomatic period is exploited before complete loss of ovary function.

Here, we combine snRNA-seq and bulk RNA-seq datasets to provide insight into the anatomy and transcriptional landscape of the developing human fetal ovary across a critical time period, uniquely doing so through a clinically-focused lens.

## Results

### The ovary undergoes marked morphological changes during fetal development

In order to characterize changes in human ovary morphology between Carnegie Stage (CS) 22/23 (8wpc) and 20wpc (Figure 1a), detailed measurements of ovary size were coupled with micro-focused computed tomography (micro-CT) analysis of anatomical location of the ovary in the developing female fetus, and with high resolution micro-CT-generated images of ovarian surface anatomy.

**Figure 1.**
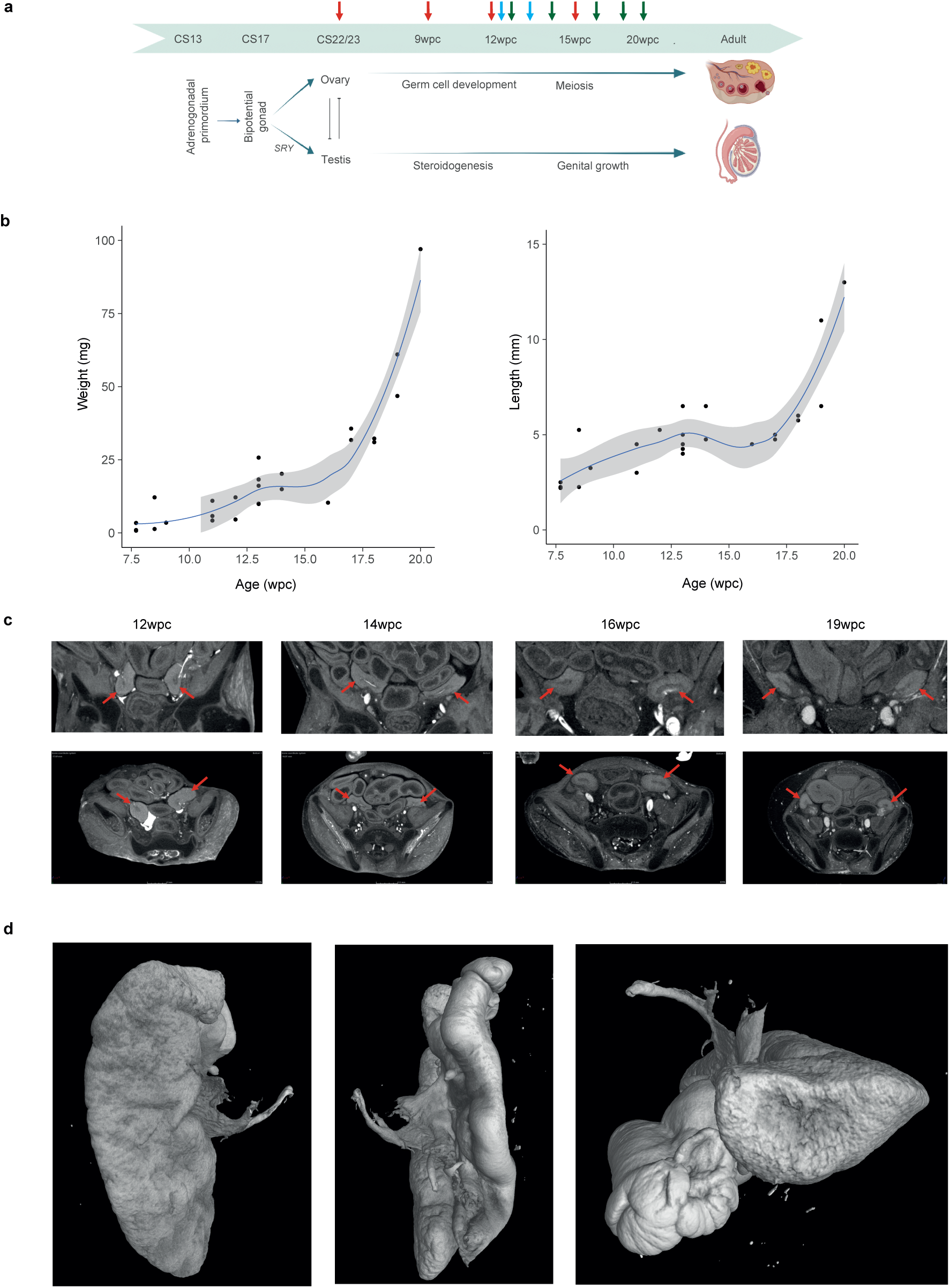
Human fetal ovary growth and morphology during development. **a** Graphic depicting the antagonistic differentiation pathways of gonads arising from the adrenogonadal primordium at approximately CS17 and resulting in the formation of the fetal ovary or testis. Red arrows indicate experimental stages for the bulk RNA sequencing study; blue arrows indicate stages included for the single-cell RNA sequencing study. CS, Carnegie stage; wpc, weeks post conception. **b** Growth curves of the ovary between CS22 and 20wpc (left panel: weight (mg); right panel: length (mm)). Data for single glands (n=27) are shown. **c** Micro-CT of the fetal ovary *in situ* at 12wpc, 14wpc, 16wpc, and 19wpc demonstrating growth and anterolateral migration of the ovary over time. Ovaries are indicated with red arrows. Upper panels show coronal views and lower panels show axial views. **d** Micro-CT images of a fetal ovary and associated Fallopian tube at 20wpc.

Fetal ovary growth curves (n=27 samples) were generated that showed substantial changes in size across this critical time period, with the most marked increase in weight and length from 17wpc (Figure 1b; Supplementary Table 1). The fetal ovary also undergoes distinct anatomical changes, as shown by coronal and axial micro-CT images of four ovaries *in situ* between 12wpc and 19wpc (Figure 1c). These images demonstrate the anterior and lateral migration of both ovaries within the abdomen in early development, from their early location on the posterior abdominal wall near the adrenogonadal primordium to a more lateral position in the developing pelvic cavity. By 20wpc, micro-CT clearly shows that the ovary has a distinct blood supply entering the hilum and is in close apposition to the developing fimbriae of the Fallopian tube, as is typical of adult anatomy (Figure 1d; Supplementary Video 1).

### The fetal ovary and testis become transcriptomically distinct with age

In order to define transcriptomic changes over time, bulk RNA-seq was undertaken using a total of 47 embryonic/fetal organs at four developmental stages (CS22-23 (7.5 to 8 wpc); 9-10wpc; 11-12wpc; 15-16wpc). This included 19 ovaries (46,XX; n=5 biological replicates at each of CS22/23, 9/10wpc, and 11/12wpc; n=4 replicates at 15/16wpc); 20 testes (46,XY; n=5 replicates at each of the four developmental stages); and eight 46,XX control tissues (two different tissue samples per development stage, including spleen, skin, kidney, muscle, stomach, lung, and pancreas) (Supplementary Figure 1). The selection of control tissue ensured a mix across endo-, meso-, and ectoderm cell lineages, and at similar stages of development.

Gene expression profiles between the three groups – ovary, testis, and control – were compared using principal component analysis (PCA) and by generating heatmaps and cluster dendrograms. Samples clearly clustered together according to developmental stage and tissue type (Figure 2a). PCA analysis demonstrated that the testis and ovary are more similar to one another than to corresponding control tissue, particularly at earlier developmental stages, and adopt a divergent transcriptomic profile with time (Figure 2a, Supplementary Figure 2).

**Figure 2.**
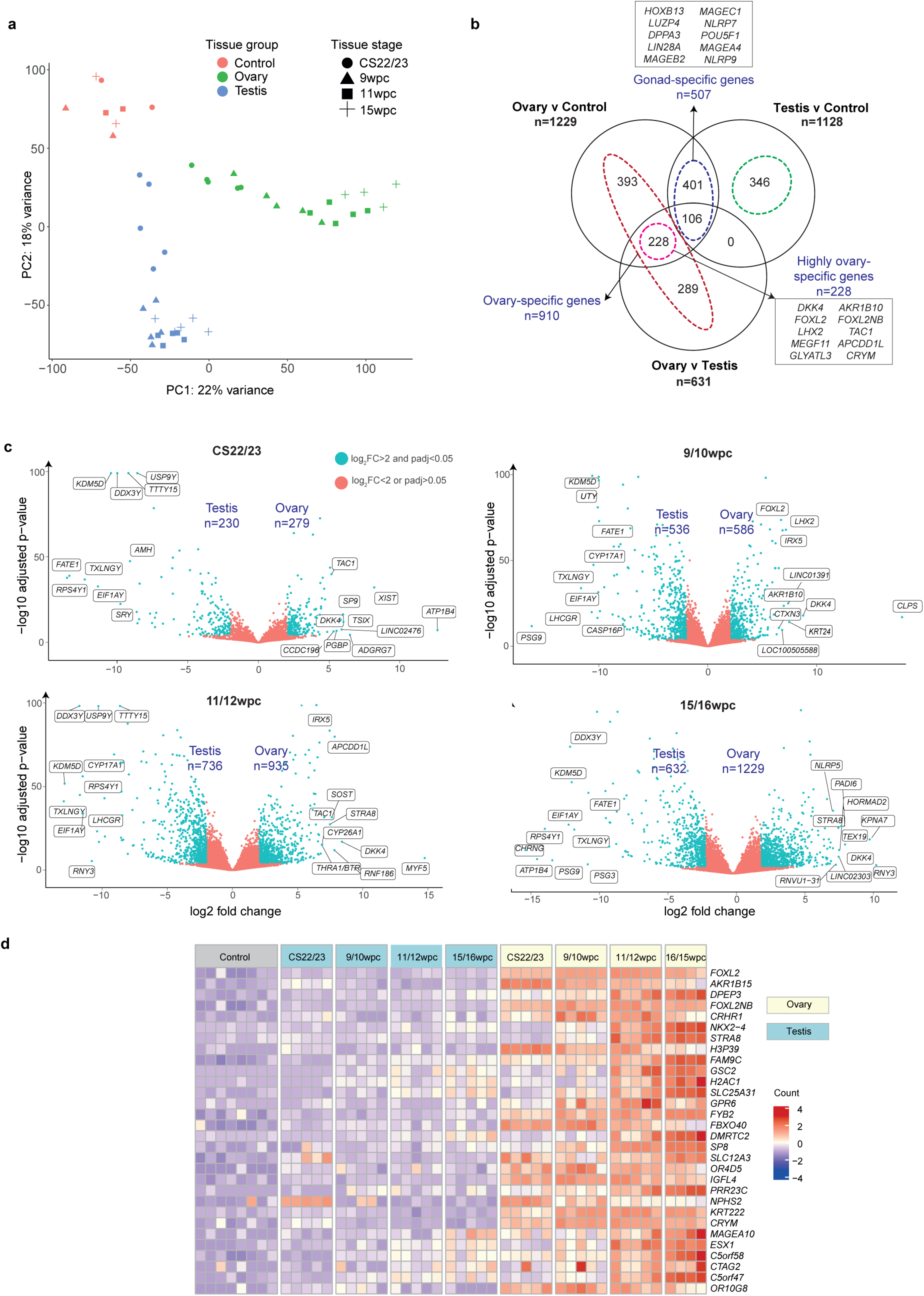
Transcriptome analysis during human fetal ovary development. **a** Principal component analysis of all 47 tissue samples included in this study (19 ovaries, 20 testes, 8 controls). The plot shows individual samples positioned according to the first and second principal components. CS, Carnegie stage; PC, principal component; wpc, weeks post conception. **b** Overlay between three differential gene expression analyses: ovary *v* control; testis *v* control; ovary *v* testis (log_2_FC>2, p.adj<0.05). The total numbers of positively differentially-expressed genes (DEGs) are shown above the Venn diagrams. Total numbers and names of the top ten genes within the ovary-specific (red), testis-specific (green), gonad-specific (violet), and highly ovary-specific (pink) DEGsets are shown within the Venn diagram. “Highly ovary-specific” refers to DEGset in both the ovary *v* control and ovary *v* testes datasets. The top ten differentially expressed genes within the gonad-specific and highly ovary-specific genesets are shown in boxes. **c** Volcano plots demonstrating four differential gene expression analyses comparing ovary *v* testis samples at CS22/23 (both n=5), 9/10wpc (both n=5), 11/12wpc (both n=5), 15/16wpc (ovary n=4, testis n=5). Each dot represents a single gene. Genes that were differentially expressed with a log_2_FC>2 and p.adj<0.05 are coloured cyan; those coloured red did not meet these significance thresholds. A max value of log_10_(p.adj) is displayed on the y-axis of the graph. Total numbers of positively and negatively DEGs are shown in violet. Gene names of the top ten positively and negatively DEGs across each developmental timepoint are displayed, based on log_2_FC. **d** Heatmap showing normalized gene expression values for the top 30 differentially expressed highly ovary-specific genes identified in (b) across the entire count matrix. Genes are ordered according to descending log_2_FC values. Samples are ordered according to tissue and stage (supervised). A color scale represents gene expression intensity (violet, lowest; yellow, neutral; red, highest). A p.adj value of <0.05 was used throughout this analysis.

Differential gene expression analyses across all developmental timepoints were performed to identify positively and negatively differentially expressed genes (DEGs) between all 1) ovary *versus (v)* testis samples; 2) ovary *v* control samples; 3) testis *v* ovary samples; and 4) testis *v* control samples (Supplementary Tables 2-5). Overlaying highly DEGs (log_2_FC>2, p.adj<0.05) within these analyses revealed ovary-specific (positively DEGs in the ovary *v* control <u>or</u> ovary *v* testis datasets); testis-specific (positively DEGs in the testis *v* control <u>or</u> testis *v* ovary datasets); and gonad-specific (positively DEGs in the ovary <u>and</u> testis within the four analyses) factors (Figure 2b). A subset of “highly ovary-specific” genes (n=228) was compiled from genes differentially expressed in both the ovary *v* control <u>and</u> ovary *v* testis analyses (Figure 2b, Supplementary Table 6). Four time-specific differential gene expression analyses were also performed between ovary *v* testis samples at CS22/23, 9/10wpc, 11/12wpc, and 15/16wpc, and between ovary samples at CS22/23 and 15/16wpc (Supplementary Tables 7-11).

Globally, the ovary and testis had distinct transcriptomic signatures with increasing divergence in gene expression patterns over time (Figure 2c, d). Indeed, more DEGs were seen in the developing ovary compared to testis, both globally and at individual developmental stages (Supplementary Table 12). Transcripts of long non-coding RNA genes (lncRNAs) were also captured. Notably, there were significantly more differentially expressed non-coding RNA transcripts in the ovary compared to both testis and control samples (log_2_FC>2, p.adj<0.05) (Supplementary Figure 3). Of the 200 non-coding transcripts positively differentially expressed in the ovary compared to either control or testis, 65.7% (n=135) were ovary-specific and 19.5% (n=39) were highly ovary-specific (Supplementary Figure 3, Supplementary Table 13).

Gene enrichment analysis of DEGs in the ovary compared to controls (n=1229; log_2_FC>2; p.adj<0.05) demonstrated an enrichment of gene ontology terms related to meiosis, gametogenesis, and/or oogenesis (n=122 genes; 9.9% of total DEGs) as well as an enrichment of genes relating to neuroactive signaling (n=55 genes; 4.5% of total; Table 1); steroid hormone metabolism (n=55 genes; 4.5% of total); and piwi RNA (piRNA) processing (n=28 genes; 2.3% of total) (Supplementary Figure 4). Enrichment analysis of DEGs in the ovary compared to testis showed a similar enrichment of genes relating to meiosis (n=61 genes; 9.7% of total DEGs) and neuroactive signalling, neurotransmitter development, and/or neural development (n=101; 16.0% of total DEGs) (Supplementary Figure 4). The highly ovary-specific geneset (n=288) was analysed separately and was found to be particularly enriched for meiosis genes (n=29; 10.1% of total geneset); G-protein coupled receptor (GPCR) signalling (n=18; 6.3% of total DEGs); neuroactive signalling (n=9; 4.1% of total DEGs); and steroid hormone biosynthesis (n=8; 2.8% of total DEGs).

**Table 1.** Neuroactive genes in the fetal ovary. The top 30 differentially expressed genes in the developing ovary (n=19) v testis (n=20) (log_2_FC<2; p.adj<0.05) with proposed functions in neurotransmission or neuroendocrine signalling on gene enrichment analysis.

| Symbol | Log2FC | p.adj | Symbol | Log2FC | p.adj |
| --- | --- | --- | --- | --- | --- |
| **LHX2** | 5.9244 | 1.13E-56 | **LINC02210** | 3.4884 | 2.68E-16 |
| **TAC1** | 5.4319 | 2.03E-42 | **GDF2** | 3.4077 | 1.65E-07 |
| **IRX3** | 5.2222 | 1.86E-39 | **WNT8B** | 3.4065 | 2.29E-17 |
| **DCT** | 5.0134 | 1.99E-19 | **GABRG1** | 3.3770 | 1.17E-36 |
| **GRPR** | 4.7377 | 1.23E-44 | **SLC34A3** | 3.3344 | 1.31E-19 |
| **GPR6** | 4.4448 | 4.30E-17 | **PRKAG3** | 3.3148 | 7.16E-09 |
| **NPY** | 3.9248 | 1.45E-26 | **NKX6-1** | 3.3131 | 2.75E-07 |
| **RASGRF1** | 3.8351 | 6.34E-50 | **GRM4** | 3.3025 | 7.74E-30 |
| **SALL1** | 3.8184 | 7.10E-14 | **NXPH4** | 3.1454 | 1.24E-25 |
| **GRM3** | 3.6915 | 1.03E-28 | **ZIC1** | 3.1045 | 5.26E-08 |
| **GRIK1** | 3.6629 | 5.56E-29 | **TACR3** | 3.1028 | 7.09E-19 |
| **RELN** | 3.6192 | 1.09E-34 | **ZIC3** | 3.0869 | 2.94E-07 |
| **GALR1** | 3.5844 | 5.00E-11 | **FOXR1** | 3.0733 | 2.88E-26 |
| **CCR8** | 3.5679 | 9.11E-24 | **GABRA4** | 3.0423 | 1.76E-12 |
| **CRHR1** | 3.4884 | 2.68E-16 | **HTR6** | 3.0277 | 1.84E-13 |

### Combining bulk RNA-seq and snRNA-seq provides novel insight into ovary development

To link the findings in bulk RNA-seq time series analysis to individual cell types, snRNA-seq was undertaken at a critical stage of ovary development: the onset of meiosis.

For this analysis, snRNA-seq profiled a total of 10,291 cells from one 46,XX 12wpc and one 46,XX 13wpc fetal ovary (quality control data: Supplementary Figure 5) and yielded a UMAP projection of the snRNA-seq transcriptome of the peri-meiotic ovary (Figure 3a). Cell annotation was assigned using established and recently identified markers in previous single-cell sequencing analyses of gonadal tissue (Figure 3b, Supplementary Table 14)^13,14^. The bulk RNA sequencing and single-nuclei data were then used to localize key DEGs of interest to cell population and developmental stage. To validate this approach, the datasets were first used to explore genes with established function in ovarian development. Germ cell markers (*DDX4, SYCP1, STRA8, MEIOC, DAZL)* and granulosa cell markers (*GATA4, KITLG, NR2F2, UPK3B, IRX3)* were expressed in the expected cell subpopulations and at predicted developmental timepoints (Figure 3b). Next, the single-nuclei data were used to explore highly differentially expressed ovary-specific genes discovered in the bulk RNA-seq dataset (log_2_FC>2.5) that had no, or limited, reported associations with ovary development. Several potentially novel patterns emerged.

**Figure 3.**
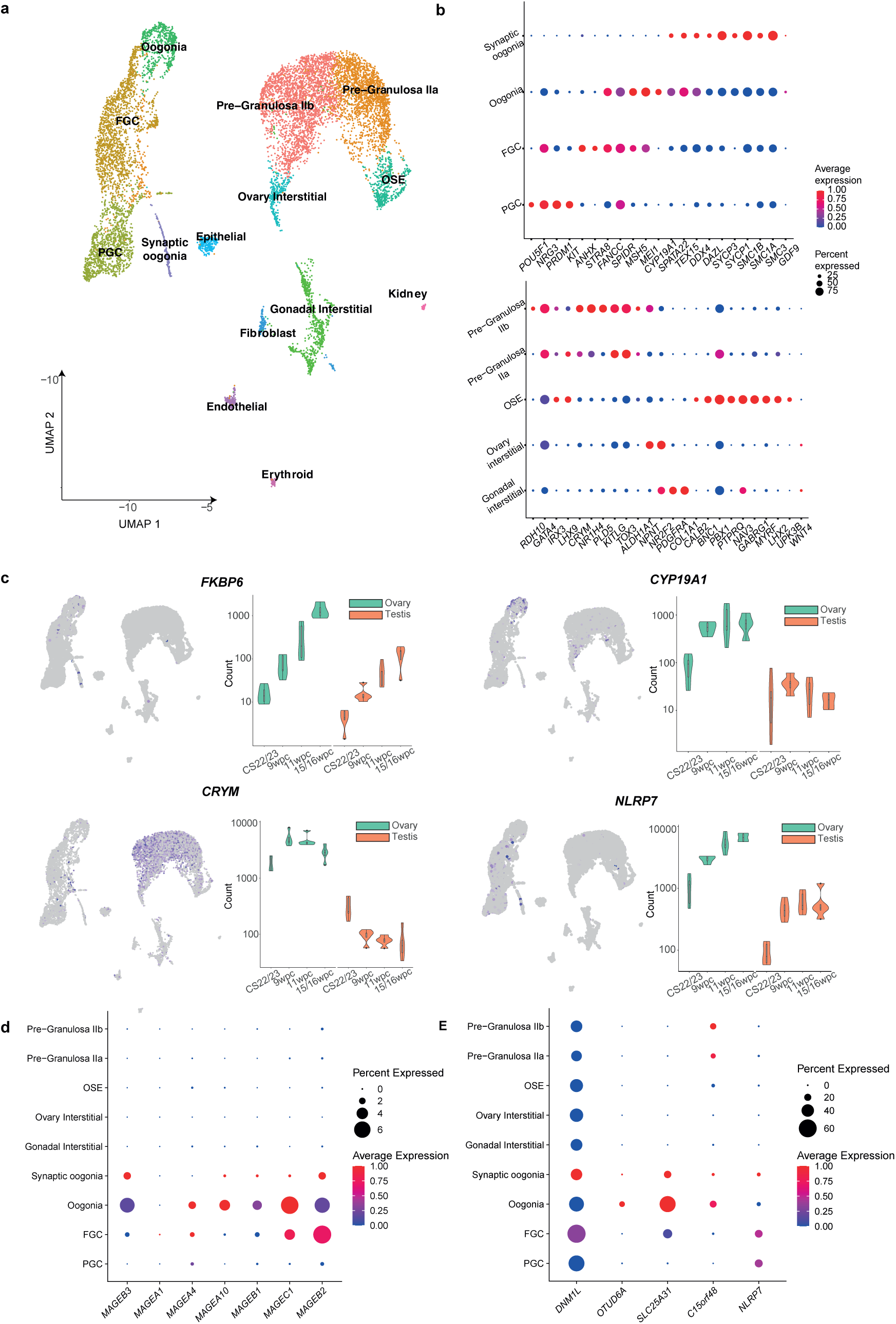
snRNAseq of two 46,XX human fetal ovaries. **a** UMAP (uniform manifold approximation and projection) of cell lineages across the merged dataset, which represents 10,291 cells from two 46,XX ovaries. Individual cell populations are annotated. FGC, fetal germ cells; OSE, ovarian surface epithelium; PGC, primordial germ cells. Markers were based on prior knowledge, markers identified in this study (e.g., *GABRG1*), and on previously published markers (Garcia-Alonso et al, *Nature,* 2022). **b** Upper panel: Dot plot showing the relative average expression and percentage of cells expressing *germ cell* markers used for snRNAseq analysis of the developing ovary. Lower panel: Dot plot showing the relative average expression and percentage of cells expressing *somatic cell* markers. FGC, fetal germ cells; OSE, ovarian surface epithelium; PGC, primordial germ cells. **c** Violin plots showing key differentially expressed genes in the fetal ovary localized to the merged 11/12wpc 46,XX snRNAseq dataset (left panel; see Figure 3a for cluster annotation) and to the bulk RNAseq data at four developmental stages. **d** Dot plot demonstrating expression within different single-cell populations of key MAGE family genes. **e** Dot plot demonstrating expression within different single-cell populations of potentially upregulated novel ovary genes, some of which are related to metabolic function or cell proliferation.

Firstly, a high number of DEGs with acknowledged roles in male gonadal and germ cell development, but with little previously described roles in the ovary, were identified. For example, *FKBP6,* the FK506 binding protein, has an essential role in male fertility and meiosis, but no acknowledged role in female gametogenesis^15,16^. *FKBP6* was highly differentially expressed in the ovary compared to controls (log_2_FC 7.67) and testis (log_2_FC 2.55) in this analysis. *FKBP6* had higher expression at female meiosis and localized to fetal germ cells (FGC) and oogonia cell clusters (Figure 3c). In the testis, *FKBP6* is required for piRNA metabolism and normal pachytene progression^17^. Here, gene expression within the developing ovary was found to be particularly enriched for piRNA-related genes. Thus, *FKBP6,* possibly within piRNA networks, may have a role in female as well as male meiosis.

*MAGE* genes (melanoma antigen genes) are members of the cancer-testis family classically associated with spermatogonia development where they protect the male germline against environmental stress^18^. *MAGE* genes have little known function in female tissues and are not expressed in the adult ovary in the Human Protein Atlas^19^. However, *MAGEA1* has been associated with proliferating germ cells in the developing ovary^20,21^. In our dataset, *MAGEA1* was differentially expressed in the ovary compared to control tissue (log_2_FC 2.77). Other *MAGE* genes were very highly differentially expressed in the ovary; for example, *MAGEB2*, *MAGEC1, MAGEA10,* and *MAGEB1*, all X chromosome genes, were among the top 50 differentially expressed genes in the ovary compared to controls (log_2_FC 9.82, log_2_FC 9.70, and log_2_FC 8.66, respectively) and *MAGEA10* was one of the most highly ovary-specific genes (Figures 2d, 3d, Supplementary Table 15). Although overall counts were modest, the expression of these genes increased at meiosis and clustered to developing germ cells in snRNA-seq data (Figure 3d). These data suggest an unexplored function for the *MAGE* gene family during germ cell development in the fetal ovary.

Highly differentially expressed genes with no known role in gonadal development were also identified. For example, *CRYM,* a highly-specific ovary gene here (ovary *v* testis log_2_FC 4.93; ovary *v* control log_2_FC 4.91), has no previously described function in ovarian development but is known to modulate thyroid hormone receptor activation^22^. Thyroid hormone has been proposed to regulate steroidogenesis in the developing granulosa cell in response to gonadotropins^23,24^; here *THRA,* and to a lesser extent *THRB*, localised to pre-granulosa cell populations (Supplementary Figure 6). In these data, *CRYM* clearly localized to the pre-granulosa cell clusters and was highly expressed from 9/10 weeks onwards (Figure 2d, 3c)^22,23^. Notably, *CYP19A1*, encoding aromatase (an enzyme essential for estrogen biosynthesis and folliculogenesis), was another highly ovary-specific gene in our dataset (ovary *v* testis log_2_FC 4.13; ovary *v* control log_2_FC 4.05). *CYP19A1* is known to be expressed from granulosa cells; interestingly, here, it was mostly expressed from oogonia cell populations (Figure 3c). This posits a potential role for crosstalk between pre-granulosa cell and oogonia populations in the earliest stages of folliculogenesis in fetal life.

Another highly expressed gene in the ovary across multiple analyses was *NLRP7,* a modulator of cellular inflammation with no recognised role in human ovary development (ovary *v* testis log_2_FC 3.16; ovary *v* control log_2_FC 9.56). Expression was most marked at meiosis and localized to all primordial germ cell (PGC), FGC, and oogonia cell clusters (Figure 3c, Figure 3e). Notably, *C15orf48,* another highly expressed ovary gene (ovary *v* testis log_2_FC 4.17; ovary *v* control log_2_FC 4.70) is also known to modulate the intracellular inflammatory response^24^. *C15orf48* expression localized to pre-granulosa cells as well as oogonia, and, like *NLRP7,* had increased expression at meiosis (Figure 3e).

Of note, there was high expression within the developing germ cell of a gene subset associated with mitochondrial metabolism. These included the highly ovary-specific genes *SLC25A31* (ovary *v* control log_2_FC 5.39; ovary *v* testis log_2_FC 3.22) and *OTUD6A* (ovary *v* control log_2_FC 7.56; ovary *v* testis log_2_FC 3.29). Mice with *Slc25a31* mutations cannot effectively utilize mitochondrial ATP and their spermatogonia undergo arrest at the leptotene stage and subsequent apoptosis^25^. In the context of human colorectal cancer, low expression of *OTUD6A* leads to suppressed mitochondrial fission and reduced cell growth of proliferating cells^26^. The expression of both *OTUD6A* and *SLC25A31* in oogonia is consistent with a role in proliferating cells (Figure 3e). *DNM1L* facilitates mitochondrial fission by increasing the expression of *OTUD6A;* notably, *DNML1L* also had marked increase at the time of meiosis in this dataset^26^

### The fetal ovary is enriched for a distinct subset of transcription factors

A focused analysis of transcription factor (TF) expression in the developing ovary was conducted to identify potentially novel regulators of gene expression (n=1,638 TFs included, 6.8% of coding genes; geneset from AnimalTFDB v4.0; accessed 15/3/23)^27^. Broadly, there was an enrichment of TFs within DEGs in the developing ovary compared to testis across most developmental stages (Supplementary Table 16). There was a similar enrichment of TFs within DEGs in the ovary compared to control (Supplementary Table 16).

The set of differentially expressed TFs in the ovary compared to testis (n=77, log_2_FC >2, padj <0.05) was investigated further to identify ovary-specific transcriptional regulation networks. Globally, most TFs belonged to Homeobox (n=31; 40.3%), zf-C2H2 (n=14; 18.2%), helix-loop-helix (bHLH) (n=7; 9.1%), or Fork-head (n=6; 7.8%) TF families. Individual stage-specific analyses demonstrated an increase of Homeobox family TFs from 9/10wpc onwards (e.g., *LHX2, IRX3, IRX5, NOBOX);* a corresponding decrease in zf-C2H2 TFs from CS22/23 (e.g., *ZNF727, ZNF729*) (Figure 4a); and a subset of TFs highly expressed at meiosis (15/16wpc) (Figure 4b). Overlaying differentially expressed TFs (log_2_FC>2, padj<0.05) from the ovary *v* control, ovary *v* testis, testis *v* control, and testis *v* ovary differential gene expression analyses yielded gonad-specific, ovary-specific, and testis-specific TFs (Figure 4c, 4d). Gene enrichment of ovary-specific TFs revealed gene ontology terms including sex differentiation, gametogenesis, central nervous system differentiation, and olfactory bulb development (Figure 4e). Several of the differentially expressed TF-encoding genes identified above have established functions in the developing ovary, such as *FOXL2, LHX8, NOBOX, LIN28A, POU5F1, FIGLA,* and *PRDM14,* and gonad-specific roles for others were described in a recent single-cell atlas^13^. Some, such as *ZIC1,* have plausible biological links to gametogenesis or gonadal development but as yet unknown roles in the developing ovary.

**Figure 4.**
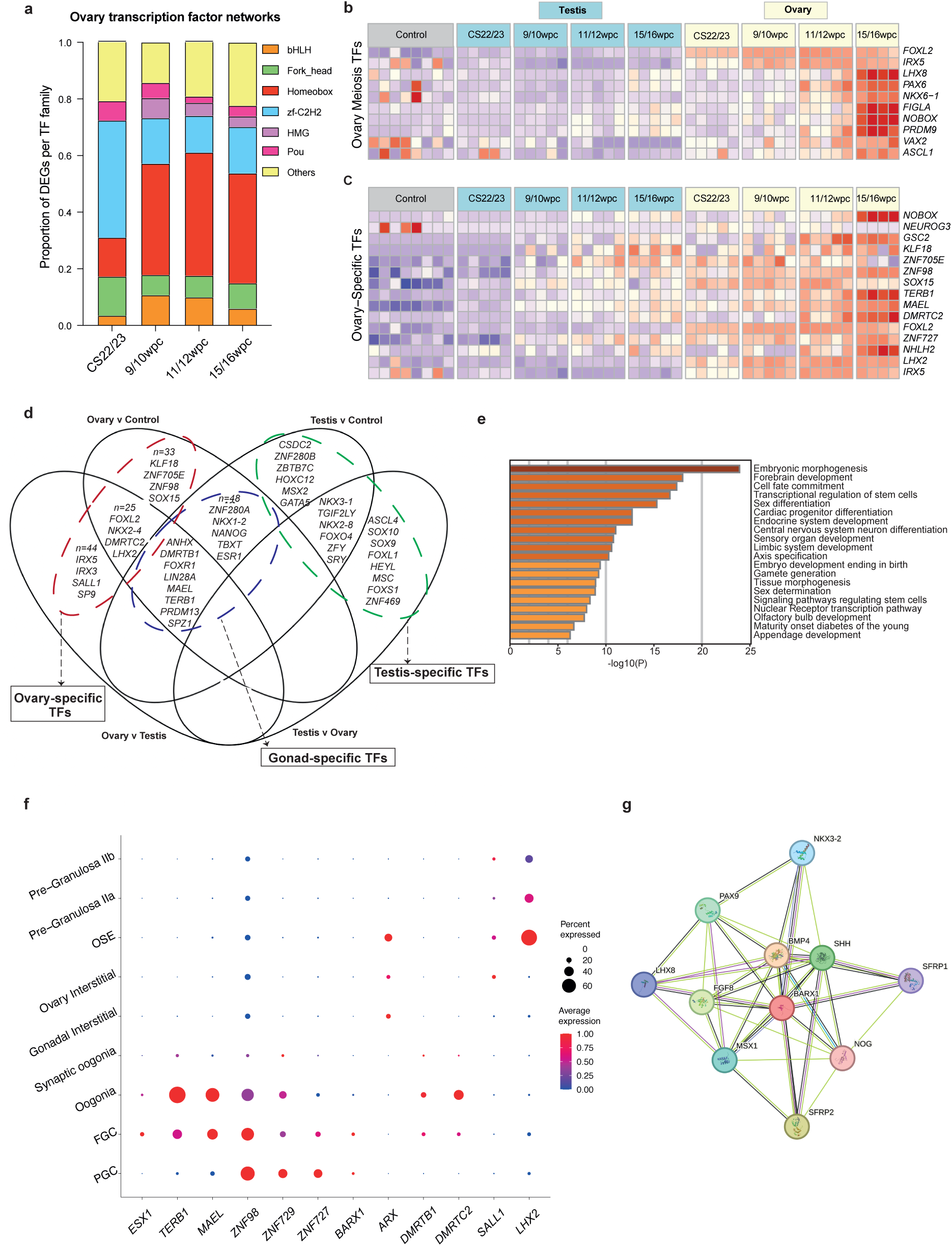
Ovary-, testis-, and gonad-specific transcription factors in early development. **a** The relative proportions of differentially expressed transcription factors (TFs) in the ovary from the most represented TF families are shown across four developmental stages (log_2_FC>2, p.adj<0.05). bHLH, Basic Helix-Loop-Helix; DEGs, differentially expressed genes; HMG, high mobility group; TF, transcription factor; zf-C2H2, Cys2–His2 zinc finger. **b** Heatmap showing top ten differentially expressed TFs at the 15/16wpc peri-meiosis stage (log_2_FC>2, p.adj<0.05). **c** Heatmap showing expression of the top 15 ovary-specific TFs across the dataset (differentially expressed in both ovary *v* testis and ovary *v* control differential gene expression analyses). **d** Venn diagram demonstrating overlap between four differential gene expression analyses (log_2_FC>2, p.adj<0.05; ovary *v* testis, ovary *v* control, testis *v* control, and testis *v* ovary) in order to identify ovary-, gonad-, and testis-specific TFs. All TFs are shown where possible, or else the total number (n) and a subset of most highly expressed in that category. **e** Gene enrichment analysis (Metascape) of ovary-specific TFs. **f** Dot plot localizing the expression of key TFs of interest to individual cell clusters on snRNAseq analysis. **g** STRING protein network analysis of BARX1.

The bulk and snRNA-seq data were used to localize the expression of TFs of interest to cell subpopulations and developmental timepoints (Figure 4c, 4f). *LHX2,* a highly ovary-specific TF (ovary *v* testis log_2_FC 5.92, ovary *v* control log_2_FC 4.86) proposed to suppress endothelial cell migration and vascularization in the developing ovary^28^, localized to pre-granulosa cells and particularly to the ovarian surface epithelium (OSE) cluster (Figure 4c, 4f)^13^. The expression of *SALL1* was higher in the ovary *v* testis (log_2_FC 3.82) and localized to the pre-granulosa and ovary interstitial populations (Figure 4f). Expression of *DMRTC2* and *DMRTB1* within developing oogonia has been previously shown in mice, macaque, and humans^8,29,30^. Here, expression of these genes localized to oogonia populations and was highest at meiosis, providing further evidence for a meiotic role in humans. *ARX,* a gene important for mammalian germ cell development and associated with X-linked lissencephaly and testis dysgenesis, was also identified as a key differentially expressed TF in this dataset and localized to gonadal interstitial and OSE populations^31^. *BARX1,* a Homeobox TF involved in Wnt signalling, has no known ovary role but was differentially expressed in the ovary compared to testis in our data (log_2_FC 2.13). Its expression increased at meiosis and localized to PGCs and FGCs. *BARX1* is predicted to interact with *LHX8* (a TF essential for oocyte development) on STRING protein network analysis, highlighting a potential role for it in human gametogenesis (Figure 4g)^32^. TFs with roles in meiotic progression were also identified: *MAEL,* a piRNA gene governing microtubule organisation within the oocyte; *TERB1,* required for telomere attachment during meiotic homologous recombination; and *ESX1*, associated with retinoic-acid responsive FGC development. These three genes are required for normal meiosis in mice and monkeys^33–36;^ in humans, *TERB1* and *ESR1* pathogenic variants lead to spermatogenic failure and male infertility^37,38^. In our dataset, *MAEL, TERB1,* and *ESX1* were strongly expressed in the ovary, localized to germ cell populations, and exhibited peri-meiotic expression (Figure 4f). Consistent with meiotic roles, *MAEL* and *TERB* peaked at 15/16wpc and clustered to later germ cell clusters (Figure 4c, 4f). *ESX1* had modest expression in FGCs, with highest expression at 11/12wpc.

### A subset of nuclear receptors is expressed in the developing ovary

Nuclear receptors (NRs) are a key subfamily of transcription factors which often have roles in gonadal development and disease (e.g., *NR5A1/*steroidogenic factor-1*, NR2F2/*COUP-TFII)^39^. Given this, the ovary *v* control, ovary *v* testis, testis *v* control, and testis *v* ovary datasets were systematically analyzed for differential expression (log_2_FC>0.5, p.adj<0.05) of the 48 known human NRs. As before, datasets were conflated to identify gonad-specific, ovary-specific, and testis-specific NRs (Figure 5a and b). This approach revealed differential expression of NRs with established gonadal roles, such as gonad-specific expression of *ESR1* (encoding the estrogen receptor-alpha), *ESR2* (estrogen receptor-beta), and *NR6A1* (encoding GCNF, germ cell nuclear factor), as well as NRs with less well-established roles in the developing ovary. For example, *NR1H4,* also known as the farnesoid X/bile acid receptor, was highly expressed at later ovary stages and, as seen previously, mostly localized to pre-granulosa IIb cells (Figure 5b)^8^. *PPARG* (peroxisome proliferator activated receptor gamma) was an ovary-specific NR with expression at all developmental timepoints localizing to germ cell populations (Figure 5b). *NR3C2,* encoding the mineralocorticoid receptor, was also identified as an ovary-specific NR with expression in both germ cell and granulosa cell lineages (Figure 5c). Variants in *NR3C2* have been linked to Type 2 diabetes, depression, hypertension, and, recently, polycystic ovarian syndrome^40,41^. However, roles in ovarian insufficiency or gonadal dysgenesis phenotypes have not been identified. *NR2E1* (encoding TLX) and *NR2E3* (encoding PNR), two NRs more typically associated with retinal development, showed ovary-specific expression that increased with age, most marked at meiosis^39^. The two key NRs previously linked to reproductive development, *NR5A1* (encoding steroidogenic factor-1) and *NR0B1* (encoding DAX-1), were both identified in the gonad *v* control genesets but with higher differential expression in the developing testis (Figure 5b; Supplementary Table 17).

**Figure 5.**
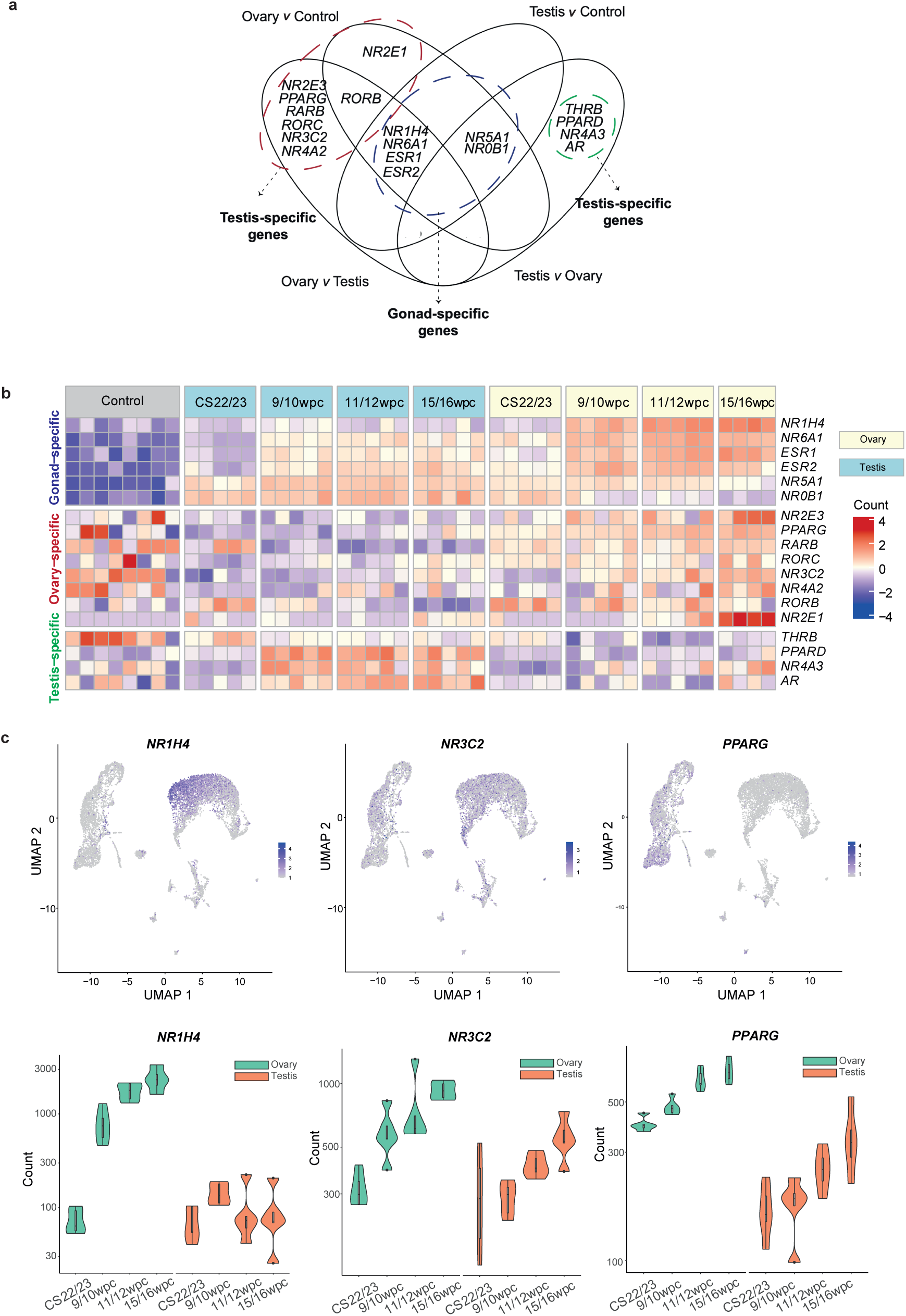
Nuclear receptor expression in the developing human fetal ovary. **a** Venn diagram showing overlapping differential gene expression analyses of nuclear receptors (NRs): ovary *v* control, ovary *v* testis, testis *v* control, testis *v* ovary (log_2_FC>0.5, p.adj<0.05). Ovary-, testis-, and gonad-specific NRs are identified. **b** Heatmap showing differentially expressed NRs across the geneset (log_2_FC>0.5, p.adj<0.05). **c** Selected highly expressed NRs localized to UMAPs from snRNAseq data (upper panels) and as violin plots over time in the bulk RNAseq data (lower panels). See Figure 3 for cluster annotation.

### Complex neuroendocrine signalling exists in the fetal ovary

An enrichment of terms relating to neurotransmitter signaling, neuroendocrine networks, and neural development was a consistent feature of genes highly expressed in the developing ovary across multiple analyses (Figure 6a, 6b, 6c; Supplementary figure 4). This finding was more marked at certain timepoints; for example, at 9/10wpc, 25.4% of all differentially-expressed genes in the ovary *v* control (log_2_FC>2; p.adj<0.05; n=149/587 genes) related to these functions (Figure 6c). Specific genes of interest emerged consistently from these analyses (Table 1). For example, *NPY* (encoding neuropeptide Y), a neuropeptide expressed by interneurons of the sympathetic nervous system, was differentially expressed in the ovary *v* testis (log_2_FC 3.92), the ovary *v* control (log_2_FC 4.07), and particularly highly expressed at earlier developmental timepoints (9/10wpc ovary *v* testis log_2_FC 4.59). Receptors *NPY1R* (Y_1_)*, NPY4R* (Y_4_), and *NPY4R2* (paralogue *NPY4R,* Y_4-2_) were also highly expressed in the developing ovary. *NPY* was highly expressed from large numbers of OSE cells as well as some pre-granulosa cell populations; *NPY4R* from small numbers of ovary interstitial, oogonia, and pre-granulosa cell populations; and *NPY4R* from small numbers of FGC populations (Figure 6d). *GABRG1, GABRA2,* and *GABRA4,* coding for three of the six alpha subunits of the GABAergic receptor, were also highly expressed alongside *NPY;* both GABAergic genes and *NPY* expression localized to OSE clusters (Figure 6a, 6b; Table 3). Genes involved in neural differentiation, such as *NAV3, IRX3, IRX5, CRHR1*, *ASCL1, TAC3R,* and *TAC1,* were also highly expressed in the early ovary (Figure 6d). Lastly, olfactory receptors, including *OR4D5* and *OR10G8,* were highly expressed in the developing ovary, localizing to subpopulations of pre-granulosa and interstitial cell populations (Figure 6d). Together, these data suggest complex, ovary-specific neuroendocrine programmes exist in the human developing fetal ovary, particularly expressed from somatic cell populations.

**Figure 6.**
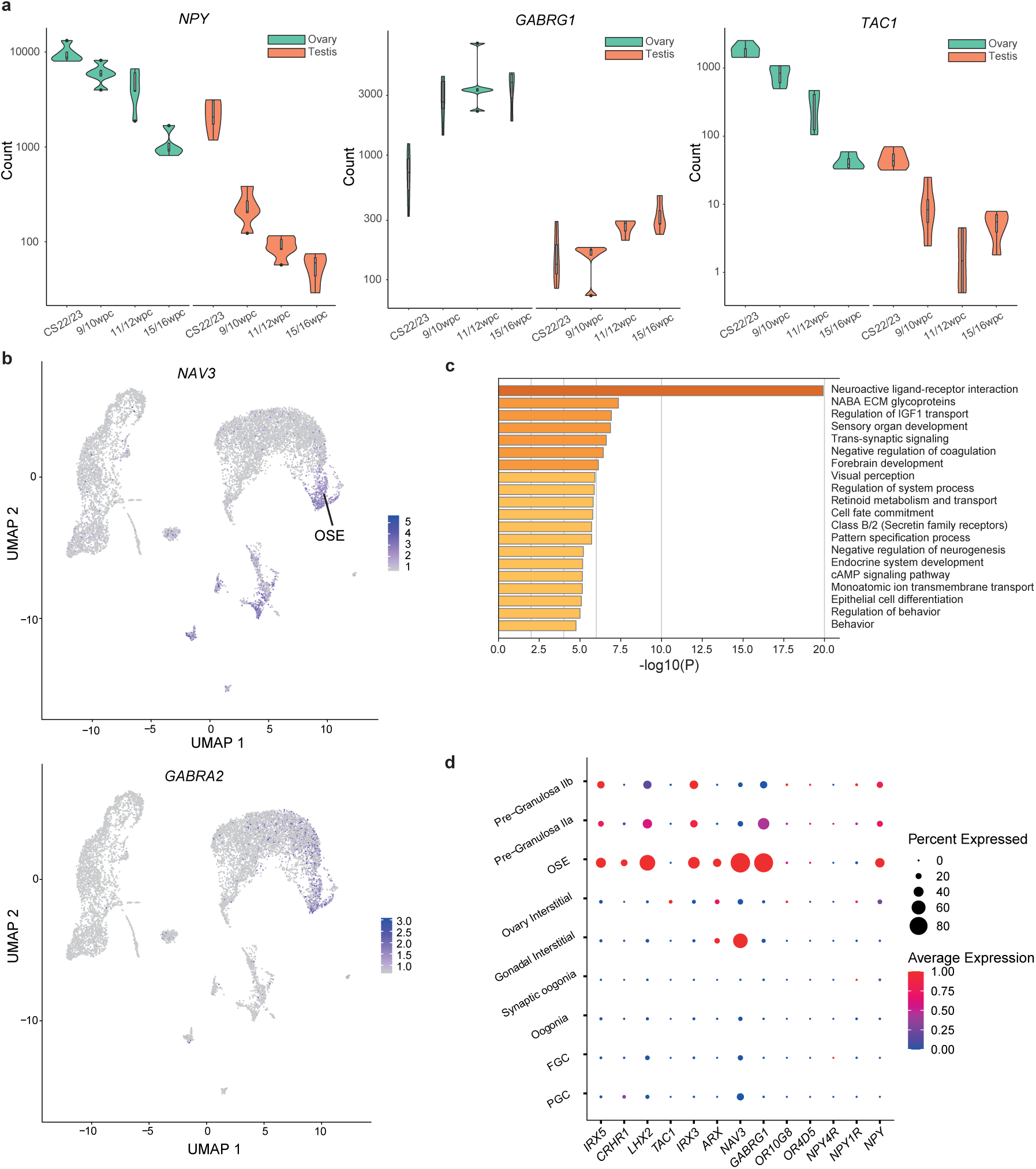
The fetal ovary is enriched for neuroendocrine and neurotransmitter genes. **a** Violin plots demonstrating the increased expression of selected neuroactive genes in the ovary at four developmental timepoints. Green, ovary; *o*range, testis. **b** UMAP representation of 46,XX ovary snRNAseq data. Expression of *NAV3* and *GABRA2* localizes to the ovarian surface epithelial (OSE) cell cluster and to interstitial cell populations. See Figure 3b for cluster annotation. **c** Gene enrichment analysis (Metascape) of differentially expressed genes (log_2_FC>2, padj<0.05) in the 9/10wpc ovary demonstrating an enrichment of terms relating to neuroendocrine and neurotransmitter signalling. **d** Dot plot demonstrating expression of selected neuroactive genes within individual cell clusters in snRNAseq analysis. FGCs, fetal germ cells; OSE, ovarian surface epithelium; PGCs, primordial germ cells.

### A focused peri-meiotic analysis reveals new meiosis candidate genes

Meiosis begins at approximately 11/12wpc and peaks at 15/16wpc. To identify genes with convincing roles in meiosis, separate analyses of differentially expressed genes in the ovary *v* testis at 15/16wpc and in the 15/16wpc ovary *v* CS22/23 ovary were conducted (Supplementary Tables 6 and 10). Gene enrichment analysis (Metascape) annotated a high proportion of differentially expressed ovary genes in either of these two analyses with meiotic gene ontology terms (n=209), including piRNA processing, meiotic cell cycle, homologous recombination and DNA repair, and cilium movement (Figure 7a). To date, several pathogenic variants in meiosis genes have been associated with the pathogenesis of primary ovarian insufficiency (POI), many related to meiosis and/or DNA repair. This was corroborated by a clear enrichment in the developing ovary of meiosis genes included on the “PanelApp” POI panel curated by Genomics England experts (v 1.67; accessed August 20th 2023; doi: https://panelapp.genomicsengland.co.uk/panels/155/) and linked to the UK 100,000 Genome Study (Figure 7b, Supplementary Table 18)^42^. In total, 25.0% (n=17) of the 68 genes on this panel were differentially expressed (log_2_FC>2, p.adj<0.05) across distinct developmental timepoints and 70.6% of these (n=13 of 17) differentially expressed PanelApp genes were included in the 209 genes annotated with meiotic terms by enrichment analysis.

**Figure 7.**
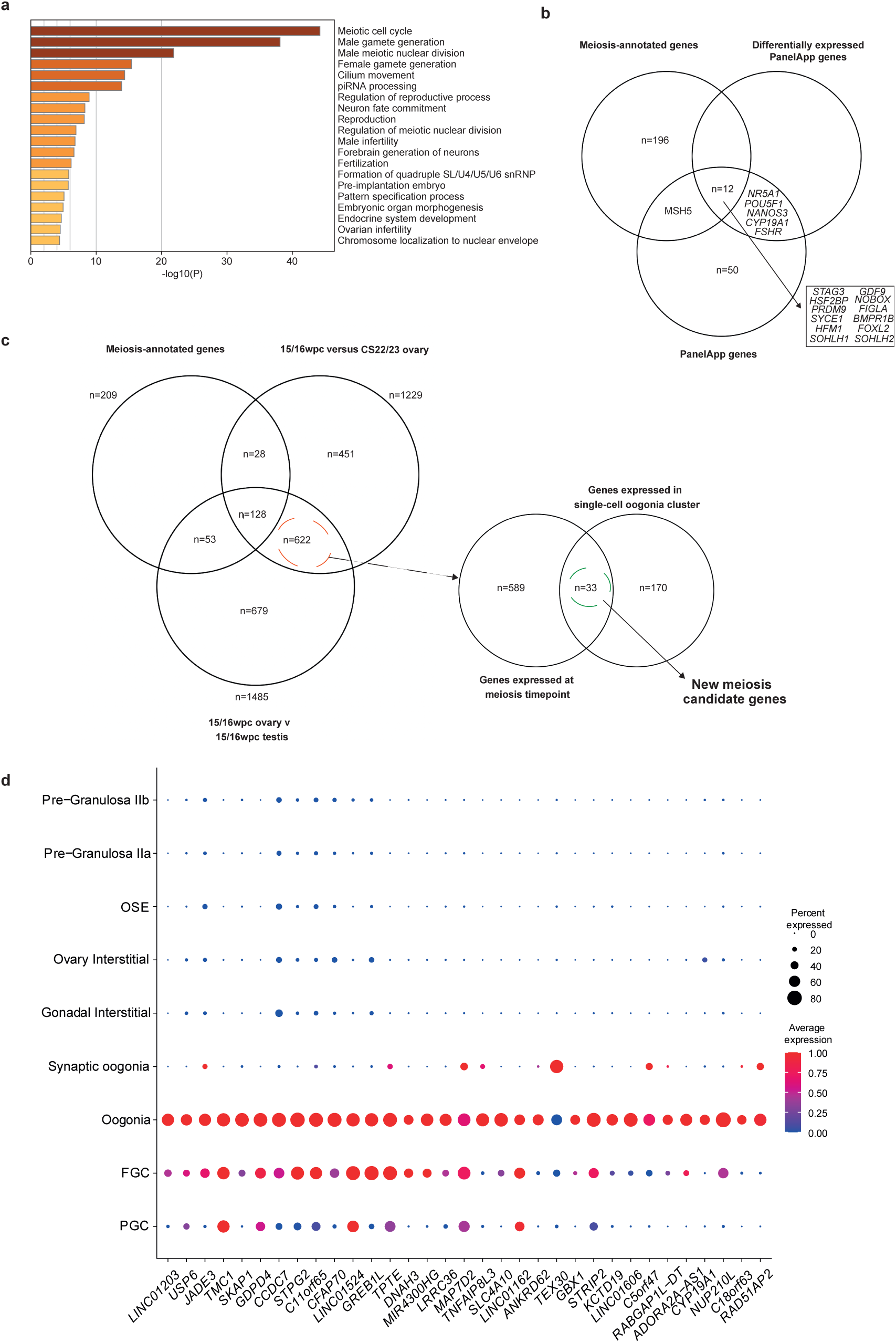
Identifying novel meiosis candidate genes. **a** Gene enrichment analysis (Metascape) of differentially expressed genes (log_2_FC>2, p.adj<0.05) at 15/16wpc, confirming an enrichment of genes related to meiosis and germ cell competence. **b** Venn diagram overlapping the PanelApp geneset; differentially expressed PanelApp genes in the ovary *v* testis (log_2_FC>2, p.adj<0.05); and meiosis-annotated genes on enrichment analysis of genes differentially expressed in the 15/16wpc ovary *v* 15/16wpc testis <u>and</u> 15/16wpc ovary *v* ovary CS22/23. **c** Genes differentially expressed in the 15/16wpc ovary *v* 15/16wpc testis and 15/16wpc ovary *v* ovary CS22/23 which were <u>not</u> annotated on enrichment analysis as meiosis genes were identified (n=622) and overlapped with markers of the oogonia cluster (n=33) to extract potentially novel candidate meiosis genes. **d** Dot plot showing expression of 33 potentially novel meiosis candidate genes across cell clusters.

We then combined transcriptomic with the above clinical data in order to try to uncover novel candidate meiosis genes. First, DEGs in both the 15/16wpc ovary *v* CS22/23 ovary and the 15/16pcw ovary *v* testis analyses that were *not* included in the list of 209 meiotic genes were identified (Figure 7c). Next, these genes were overlaid with differentially expressed genes from the oogonia cluster in the snRNAseq dataset. This approach led to the identification of 33 novel and convincing candidate genes for roles in meiosis and, accordingly, potential genetic causes of POI (Figure 7c, Table 2). Each of these candidate genes had increased expression at meiosis in the bulk RNAseq data and localized to meiotic germ cell clusters in the snRNA-seq data (Figure 7d).

**Table 2.** Novel meiosis candidate genes. Novel meiosis genes identified in this study are shown. Genes differentially expressed at meiosis-specific timepoints (15/16wpc ovary *v* testis; 15/16wpc ovary *v* CS22/23 ovary) that were not meiosis-annotated on enrichment analysis were identified and compared with oogonia specific markers from snRNA-seq analysis. Overlapping genes were considered novel candidate meiosis genes.

| *Symbol* | *Gene name* | *Proposed function* |
| --- | --- | --- |
| **TNFAIP8L3** | TNF alpha-induced protein 8-like 3 | Positively regulates phosphoinositide 3-kinase activity, which is involved in cell growth, survival, sensitivity to IGF-1, and energy restriction |
| **SLC4A10** | Sodium-driven chloride bicarbonate exchanger | Regulates the pH of neurons, choroid plexus, brain extracellular fluid |
| **STRIP2** | Striatin interacting protein 2 | Involved in cell migration, cytoskeleton organization, cell shape regulation of cell shape |
| **NUP210L** | Nucleoporin 210 like | Involved in Sertoli cell development and spermatid development |
| **SKAP1** | Src kinase associated phosphoprotein 1 | Positively regulates T-cell receptor signaling by enhancing the MAP kinase pathway |
| **RAD51AP2** | RAD51 associated protein 2 | Involved in double-strand break repair during homologous recombination |
| **MIR4300HG** | Non-coding RNA gen | Variants linked to adolescent idiopathic scoliosis; no ovary link |
| **LINC01606** | Non-coding lncRNA gene | Possibly associates with miR-423-5p to regulate Wnt/β-catenin signalling |
| **GDPD4** | Glycerophosphodiester phosphodiesterase domain-containing protein 4 | Involved in lipid metabolic processes |
| **CYP19A1** | Cytochrome P450 family 19 subfamily A member 1 | Cytochrome P450 superfamily, steroid hormone, catalyses oestradiol and aromatase synthesis |
| **TPTE** | Transmembrane phosphatase with tensin homology | Encodes a PTEN-related tyrosine phosphatase possibly involved in signal transduction pathways modulating spermatogenic function |
| **LINC01203** | RNA Gene, and is affiliated with the lncRNA class | No known function but highly expressed in testis alone in the Lncipedia |
| **GREB1L** | GREB1 like retinoic acid receptor coactivator | Causes autosomal dominant non-syndromic deafness and renal agenesis; causes CAKUT (congenital abnormalities of the kidneys and urinary tract); increases PAX2 and PTH1R; has been associated in case reports with streak ovaries and sensorineural hearing loss |
| **LRRC36** | Leucine-rich repeat-containing protein 36 | Localises to centrosome; paralogue of centrosomal protein CEP72 which recruits key centrosomal proteins to the centrosome and forms a bipolar spindle via microtubule-nucleation activity |
| **ADORA2A-AS1** | Non-coding RNA gene | Increases proliferation and is anti-apoptotic in CML context |
| **CFAP70** | Cilia and flagella associated protein 70 | Involved in cilium assembly and cilium movement; part of outer dynein arm; causes male infertility due to asthenoteratozoospermia |
| **TMC1** | Transmembrane channel like 1 | Required for cochlear hair cell function; associated with deafness; associated with ovary function in insects |
| **STPG2** | Sperm Tail PG-Rich Repeat Containing 2 | Expressed in human testis and brain; paralogue of STPG1 which regulates mitochondrial membrane permeability and modulates apoptosis |
| **LINC01524** | Long non-coding RNA | No known function |
| **C5orf47** | Chromosome 5 open reading frame 47 | No known function |
| **LINC01162** | Long non-coding RNA | No known function |
| **KCTD19** | Potassium channel tetramerization domain containing 19 | Interacts with ZFP541 and HDAC1 to play an indispensable role in pachytene meiotic progression in male mice |
| **GBX1** | Gastrulation brain homeobox 1 | Regulates nervous system development and RNA polymerase II transcription; homeobox marker preferentially expressed by GV oocytes in humans |
| **DNAH3** | Dynein axonemal heavy chain 3 | Dynein family gene encoding for microtubule-associated motor protein complexes |
| **MAP7D2** | MAP7 domain containing 2 | Involved in microtubule cytoskeleton organization |
| **JADE3** | Jade family PHD finger 3 | Possibly involved in histone H4 acetylation |
| **C11orf65** | Chromosome 11 open reading frame 65 | Negatively regulates mitochondrial fission and protein targeting to mitochondria |
| **CCDC7** | Coiled-coil domain containing 7 | Suppresses cell proliferation via the JAK-STAT3 pathway |
| **RABGAP1L-DT** | Long non-coding RNA | No known function |
| **C18orf63** | Chromosome 18 open reading frame 63 | Expression enriched in oocytes in chickens and humans |
| **USP6** | Ubiquitin specific peptidase 6 | Involved in protein deubiquitination; deubiquitinates H2A; suppresses double-stranded DNA break activity |
| **ANKRD62** | Ankyrin repeat domain 62 | Highly expressed in testis alone in Human Protein Atlas |
| **TEX30** | Testis-expressed protein 30 | One of the male germ cell-specific testis-expressed TEX gene family |

## Discussion

We present an integrated multimodal study of human fetal ovary development during a critical early stage of growth and differentiation, extending from CS22/23 (8wpc) up until the second trimester. We harness complementary multiomic and imaging approaches with the aim of defining the anatomical and molecular landscape during this critical period and contextualise our findings clinically.

Using a novel approach of micro-CT, we have demonstrated, three-dimensionally, the significant morphological change occurring within the fetal ovary within a relatively narrow time window. We show the antero-lateral movement of ovaries in the fetal abdomen throughout the second trimester and, using high resolution surface micro-CT imaging, demonstrate that by 20wpc the human ovary has a complex structure with hilar vascularization and close apposition to the fimbriae of the oviduct (Fallopian tube), which has already adopted mature anatomical features by this stage. We show marked growth and weight of the developing ovary from 16wpc onwards, corresponding with the massive germ cell expansion and meiosis occurring at this time. We advance our understanding of germ cell development, an ovary developmental timepoint essential for its later maintenance and function: by 20-22wpc, the ovaries of 46,XX fetuses have the maximum number of oogonia (approximately 7,000,000) they will hold during their entire reproductive lifetime, representative of their lifetime ovarian reserve. Disruption of this critical timepoint results in insufficient germ cell expansion and/or accelerated apoptosis of this ovarian reserve, resulting in the fewer germ cells at birth, smaller ovaries macroscopically, and the clinical phenotype of Primary Ovarian Insufficiency (POI).

We then used a series of transcriptomic approaches to show that the fetal ovary has a clear transcriptomic blueprint across all developmental timepoints defined by thousands of differentially expressed protein-coding genes as well as non-coding transcripts. Indeed, the fetal ovary has more differentially expressed genes at the developmental timepoints studied than the testes, firmly rejecting the concept of ovary development being a “default” pathway. Genes known to prevent testis development, such as *FOXL2,* were highly expressed, confirming testis antagonism as a major pathway in ovary development^43,44^. However, we demonstrate that the fetal ovary has important and unique features beyond just testis antagonism, including anatomical and vascular organization, which are linked temporally to the key biological ovary developmental processes of germ cell expansion, meiosis onset, and meiotic arrest.

Integrating bulk- and snRNAseq throughout this work allowed us to uncover several unique features of the developing ovary, some novel, and some re-enforcing previous knowledge. Genetic networks relating to meiosis were highly expressed with a particular enrichment of genes related to DNA repair. This is not surprising, given the known role of DNA repair genes during homologous recombination at meiosis I, but does highlight the vulnerability of the growing follicle pool to DNA damage and the implications of this for transgenerational epigenetics. In keeping with this is, a proportion of POI is proposed to arise from environmental toxin exposure, and pathogenic variants in DNA repair genes independently give rise to POI phenotypes – such as *YTHDC2* and *ZSWIM7,* which we recently described^45,46^. We then combined transcriptomic with clinical data (the PanelApp gene list) to reveal novel meiosis-regulating candidate genes, providing a catalogue of candidate genes for POI going forward.

We also elucidate a previously undescribed complex network of neuroendocrine signalling, mitochondrial pathways, and energy metabolism within the fetal ovary that begins as early in ovary development as CS22. *NPY* is highly expressed in the developing ovary, a gene which is classically expressed in the arcuate nucleus (ARC) of the hypothalamus where it dampens inhibitory GABAergic activity (modulated by *GABRG1,* also highly expressed in the developing ovary). NPY has known neuroendocrine functions, including food intake regulation and energy storage, but a potential role within a neuroendocrine signalling pathway in the developing ovary has not been described. Other neuroactive genes with roles in appetite and fat regulation in adult life – *IRX3, IRX5, NHLH2* – were also highly expressed; *IRX3* and *IRX5* have postulated roles in obesity regulation as well as ovary folliculogenesis^47–49^, and mice require Nhlh2 for meiosis and fertility as well as for energy metabolism^50^. *PPARG,* a top differentially expressed nuclear receptor (NR) in the ovary compared to testis, regulates immune cell activation as well as adipogenesis and has been detected in the adult human ovary previously, specifically within granulosa cells^51^. Excessive intra-ovarian PPARG production within granulosa cells disrupts steroidogenesis and contributes to PCOS, obesity, and insulin resistance^51^. However, we localise fetal *PPARG* expression to germ cell populations, suggesting a role in the developing ovary other than steroidogenesis – perhaps relating to energy homeostasis within germ cells or to inflammation regulation (specifically macrophage activity) within the ovary^52,53^. The high expression of Nod-like receptors (*NLRP7, NLRP4, NLRP14)* within germ cell populations in our data also aligns with the concept of a role for “inflammasome” regulation in the developing ovary. The clinical phenotypes of obesity, central hypogonadotropic hypogonadism (CHH), and PCOS have previously linked fertility with energy metabolism and its neuroendocrine control – for example, excessive GABAergic activation and disrupted weight homeostasis are implicated in the pathogenesis of PCOS^54^. Our data introduces the paradigm that these pathways are required for within the ovary from the earliest points in its development in fetal life.

In addition to those with postulated neuroendocrine function, other neuroactive genes were highly expressed in the ovary from the CS22/23 timepoint. There was a marked expression of genes involved in sympathetic nervous system development, such as *ASCL1,* which interestingly localised to the ovarian surface epithelium (OSE) clusters – cells on the epithelial surface of the ovary – indicating an as yet unexplored role of sympathetic innervation to the ovarian capsule required for ovarian development. Other neurotransmitters, such as *NAV3,* also localised to OSEs; notably, epitranscriptomic downregulation of this gene has been implicated in chemotherapy resistance in ovarian cancer, suggesting a possible role of these primitive neuroactive pathways within oncogenesis in later life^55^. There was also a striking enrichment of genes related to sensorineural function, including olfactory neurons (*OR4D5*) and retinal nuclear receptor encoding genes (*TLX*, *PNR*), as well as upregulation of other neurotransmitters (*TAC1, TAC3R*). Disruption of these pathways have been previously linked to CHH (olfactory dysfunction (anosmia) and hypogonadotropic hypogonadism in Kallman Syndrome; *TAC3R* variants in CHH^56^) but not to ovary insufficiency phenotypes. Expression of olfactory receptors outside of olfactory sensory neurons is increasingly recognized and olfactory receptors have been previously identified as highly expressed in the fetal ovary^6^. Postulated non-olfactory roles for these receptors include cell-cell recognition, migration, proliferation, and apoptosis^57^, although any potential role for olfactory receptors and their respective ligands in human ovary development remains unexplored. Taken together, we propose that the fetal ovary has specific neuroendocrine axes and energy homeostasis programmes which are required for its development and lay down the infrastructure required for ovarian function much later in adult life.

This study has certain limitations, including the relatively few samples included for snRNA-seq which necessitated a focus on the key meiotic timepoint of 12wpc. Bulk RNA-seq has inherent limitations, including its propensity to overlook small but important signals from smaller cell populations, which was mitigated by cross-referencing with snRNA-seq data and by designing a well-powered study with appropriate control samples. Key strengths of this study were its integrated investigative approach and its focus on clinically relevant biological timepoints. We have uncovered several novel candidate genes, genetic pathways, and signaling networks with roles in ovary development which remain unexplored and require interrogation in future studies. We highlight the clinical relevance of these findings, which can potentially be manipulated for therapeutic benefit. We show that the fetal ovary has a remarkably complex genetic programme which underlies a developmental trajectory characterized by critical biological processes which are essential for ovarian function, and vulnerable to disruption during later life.

## Methods

### Tissue samples

Human embryonic and fetal samples used for these studies were obtained from the Human Developmental Biology Resource (HDBR). The HDBR is a Medical Research Council (MRC)/Wellcome Trust-funded tissue bank regulated by the Human Tissue Authority (www.hdbr.org). Samples were collected with appropriate maternal written consent and with ethical approval from the NRES London-Fulham Ethics Committee (08/H0712/34+5, 18/LO/0822,) and Newcastle Ethics Committee (08/H0906/21+5, 18/NE/0290). This specific work has study approval from the HDBR (project numbers 200408; 200481; 200581). All researchers involved in this study underwent appropriate Human Tissue Act training. Ovaries, testes and relevant 46,XX control tissues were identified and isolated by blunt dissection. The age of embryonic and fetal tissue was calculated by experienced HDBR researchers (London and Newcastle) using published staging guidelines, such as Carnegie staging for embryos up to 8wpc and foot length and knee-heel length in relation to standard growth data for older fetuses. Samples were karyotyped by G-banding or quantitative PCR (chromosomes 13, 16, 18, 22, X, Y) to ascertain the sex of the embryo or fetus. Samples were stored at -70°C or fixative (10% formalin or 4% paraformaldehyde) prior to use.

### Fetal ovary measurement

Fetal ovaries between CS23 and 20wpc were obtained from HDBR and fixed in 4% PFA. Ovary dimensions were measured to the nearest mm. Ovary weights were measured using an analytical balance (Pioneer PX, Ohaus).

### Micro-focus computed tomography (micro-CT)

#### Micro-CT of fetal ovaries in situ

Human fetuses (14-21 wpc, undergoing micro-CT post-mortem imaging procedures) were prepared for imaging by immersing in 2.5% potassium tri-iodide for between 4 and 16 days dependent on weight, and dried and positioned for scanning, as described previously.^58^ Micro-CT scans were carried out using a Nikon XTH225 ST or Med-X scanner (Nikon Metrology, Tring, UK) with the following factors optimized for each fetus, depending on size: Tungsten target, X-ray energy 130 kV, current 134 - 239 µA (power 17 – 31 Watts), exposure time 250 ms, two frames per 3141 projections, detector gain 24 dB, scan duration 26 minutes. Modified Feldkamp filtered back projection algorithms were used for reconstructions within proprietary software (CTPro3D; Nikon Metrology) and post-processed was undertaken using VG StudioMAX (Volume Graphics GmbH, Heidelberg, Germany) to create both axial and coronal images at 30 - 54 µm isotropic voxel sizes.

#### Micro-CT of fetal ovary surface morphology

Surface scanning was undertaken of a fetal ovary (20wpc) and attached Fallopian tube stored in 10% formalin at 4°C. Iodination was performed by immersing the sample in 1.25% potassium tri-iodide (I_2_KI) for two days, then rinsing and drying it. The sample was then wrapped in parafilm and secured to a carbon-fibre plate to eliminate any movement or desiccation during the scan. Micro-CT scans were carried out using a Nikon XTH225 ST scanner (Nikon Metrology, Tring, UK) with the following factors: Tungsten target, X-ray energy 100-110 kV, current 46-60 µA (power 6.6 Watts), exposure time 1420-2000 ms, one frame per 3141 projections, detector gain 24 dB, scan duration 75 minutes. Modified Feldkamp filtered back projection algorithms were used for reconstructions within proprietary software (CTPro3D; Nikon Metrology) and post-processed was undertaken using VG StudioMAX (Volume Graphics GmbH, Heidelberg, Germany) to create the images at 3.78-4.77 µm isotropic voxel sizes.

### Bulk RNA-sequencing

#### RNA extraction

RNA was extracted using the AllPrep DNA/RNA Mini Kit (QIAGEN N.V.) in accordance with the manufacturer’s protocol. In brief, tissues frozen at -70 degrees were homogenized using an electronic pestle (Kimble, New Jersey) and the lysate was transferred to an AllPrep DNA spin column placed in a 2ml collection tube. This was centrifuged for 30 seconds at 8000g and 250μl 100% ethanol was added to the flow-through and pipette mixed. Following this, 700μl of the sample was transferred to a RNeasy spin column and centrifuged for 15 seconds at 8000g. The flow-through was transferred to a 2ml collection tube and 700μl of Buffer RW1 added. A 15 second centrifugation followed at 8000g to wash the spin column membrane. Next, 500μl of Buffer RPE was added to the RNeasy spin column and centrifuged for 15 seconds. The flow-through was discarded and 500μl of Buffer RPE again added followed by a longer two-minute centrifugation at 8000g. Extracted RNA was diluted in 30μL of nuclease-free water applied directly to the spin column membrane. RNA concentration and 260/280 ratios were measured using a NanoDrop ND-1000 spectrophotometer (NanoDrop Technologies, Witec, Littau, Switzerland). The minimum amount of RNA required for sequencing was 50ng with a 260/280 ratio of >2.0. RNA integrity number (RIN) was assessed using an Agilent Bioanalyser (Agilent, Santa Clara, CA); all samples had a RIN >7.

#### Library preparation and RNA sequencing

Library preparations were carried out on a Hamilton StarLet (Hamilton, Reno, NV) robotic platform and library qualitative checks performed using the Tapestation 4200 platform (Agilent, California, USA). Libraries were prepared using the KAPA RNA HyperPrep Kit and sequenced on an Illumina HiSeq® 4000 sequencer (Illumina, San Diego, CA) at a minimum of 25 million paired end reads (75bp) per sample.

#### Bioinformatic analysis

Fastq reads underwent QC (FastQC, Babraham Bioinformatics) and were aligned against the genome (GRCh38, h38 to generate BAM files (STAR 2.7)^59^. featureCounts (Subread package, v2.0.2), DESeq2 (v1.28.1), and degPatterns (DEGreport package, v1.24.1) were used for gene expression quantification, differential gene expression, and detection of expression patterns, respectively^60,61^. The cut-off adjusted p-values and log_2_fold changes were 0.05 and 1, 1.5, or 2, respectively. Metascape was used for gene annotation and pathway enrichment analysis^59^. The ComplexHeatmap package in R was used to generate heatmaps representing differentially expressed genes^62^. All R packages and versions used in the bioinformatic analysis pipeline are listed in Supplementary Methods 1.

### Single-nuclei RNA-sequencing (snRNA-seq)

#### Samples

Two 46,XX fetal ovaries (12wpc and 13wpc) were obtained from the HDBR, following ethical approval and consent as described above. The karyotype of the tissue was confirmed on extracted DNA from skin biopsy and array. Samples were stored at - 70°C prior to use.

#### Single-nuclei dissociation

A previously published protocol was used to dissociate tissues into single-nuclei suspensions (Martelotto et al, Broad Institute (dx.doi.org/10.17504/protocols.io.bw6qphdw). All samples and reagents were kept on wet ice or at 4°C throughout. In brief, tissue samples were diced using a scalpel into approximately 1mm^3^ pieces. Samples were then dissociated in 300μL of Salty Ez10 Lysis Buffer supplemented with 0.2-0.5U/μL of RNase inhibitor (Sigma-Aldrich) using a 2ml Kimble douncer (Sigma-Aldrich) (10 strokes with loose pestle and 10 strokes with tight pestle) to achieve single nuclei suspensions. An extra 700μl of chilled Salty Ez10 Lysis Buffer supplemented with 0.2-0.5U/μl of RNase inhibitor was added to the samples and gently pipette mixed. Samples were incubated on ice for five minutes and slowly pipette mixed on 2-3 occasions during incubation. A MACS 70μm filter (Miltenyi Biotec) was placed in a 50ml Falcon tube on ice. The single nuclei suspension was transferred to the filter and then the filtrate transferred to a 1.5ml LoBind tube (Eppendorf). Nuclei were centrifuged at four degrees at 500g for five minutes. The supernatant was removed to leave a pellet. Salty Ez10 Lysis Buffer was added (1ml) and the pellet resuspended gently before a further five-minute incubation on ice and five-minute centrifugation as before. The supernatant was removed, 500μl of Wash and Resuspension Buffer 2 (WRB2) added without disturbing the pellet, and the sample left to sit on ice for five minutes before being gently resuspended. Samples were counted using the Luna-FL™ Dual Fluorescence Cell Counter (Logos Biosystems) and Acridine Orange/Propidium Iodide (AO/PI) Cell Viability Kit Counter (Logos Biosystems) (9μl of sample with 1μl of Acridine Orange). Both samples were within the ideal concentration of cells per μl (ideal range 700-1200cells/μl; 46,XX: 12wpc 1720cells/μl; 46,XX 13wpc: 1170cells/μl). Components for the two buffers, made up to a 100ml stock, are shown in Supplementary Methods 2.

#### Gel bead in EMulsion (GEM) Generation and Barcoding

Libraries from single nuclei suspensions were processed using the 10X Chromium Single Cell 3’ kit as per protocol (10X Genomics, Pleasonton, CA). In brief, MasterMix was prepared on ice (1X: 18.8μl RT Reagent B, 2.4μl Template Switch Oligo, 2.0μl Reducing Agent B, 8.7μl RT Enzyme C). Following this, 31.8μl of MasterMix was added to an 8-tube PCR strip on ice. The Chromium Next Gel bead in EMulsion (GEM) Chip G was assembled as per instructions. Based on total cells/μl in each sample, the 10X Genomics Cell Suspension Volume Calculator table was used to calculate the ratio of nuclease-free water: single nuclei suspension to add to row 1 of the chip. Gel Beads were dispensed into the row 2. Partitioning Oil (45μl per sample) was dispensed into row 3. 10,000 cells per sample were loaded onto the Chromium Next GEM Chip G (10X Genomics). The Chromium Controller was then run as per protocol for 18 minutes to form GEMs: a single nucleus, a single Gel Bead, and reagents.

#### Cleanup and cDNA amplification

After, 100μl of GEMs were aspirated and incubated in a thermal cycler for 55 minutes as per protocol (Bio-Rad C1000 Touch). Recovery Agent (125μl) was added to each sample at room temperature and left to incubate for two minutes resulting in a biphasic mixture of Recovery Agent and Partitioning Oil. From this mixture, 125μl was discarded from the bottom of the tube. Dynabeads Cleanup Mix was then prepared (1X: 182μl Cleanup Buffer, 8μl DynaBeads MyOne SILANE, 5μl Reducing Agent B, 5μl Nuclease-Free Water) and 200μl of this solution added to each sample and pipette mixed ten times followed by a ten-minute incubation. The samples were then placed on a magnetic separator (10X Genomics, Chromium™ Controller Accessory Kit) until the solution cleared. Supernatant was removed and 300μl of ethanol added to the pellet while on the magnet. After 30 seconds, this ethanol was removed and the sample centrifuged briefly and air dried for one minute. Elution Solution 1 (35.5μl) was then added to the sample, pipette mixed, and incubated for two minutes at room temperature. The solution (35μl) was then transferred to a new PCR strip for cDNA amplification. Here, 65μl of cDNA Amplification Reaction Mix was added to 35μl of the sample and incubated inside the thermal cycler for 45 minutes as per protocol.

#### Library preparation using the 10X Genomics platform

A 3’ gene expression library was then constructed from the purified cDNA. Fragmentation, end repair, and A-tailing followed standard 10X Genomics workflow: 10μl of purified cDNA, 25μl of Buffer EB (1X: 5 μl Fragmentation Buffer, 10 μl Fragmentation Enzyme), and 15μl of Fragmentation Mix were added to a PCR tube strip and mixed. The sample was then transferred to a pre-cooled thermal cycler (four degrees) and run as per protocol for 35 minutes. Double-sided size selection followed: 30μl of SPRIselect (0.6X) reagent was added to each sample, mixed, and incubated for five minutes at room temperature. Samples were placed on the magnet until the solution cleared. Then, 75μl of the supernatant was transferred to a new PCR strip and 10μl SPRIselect (0.8X) added to each sample. Samples were incubated at room temperature for five minutes and then again placed on the magnet until the solution cleared. The supernatant was removed (80μl) and 125μl of ethanol added to the pellet. After 30 seconds, ethanol was removed. These size selection steps were repeated twice. Samples were then incubated at room temperature and 50μl transferred to a new PCR tube strip for adaptor ligation, which involved adding 50μl of Adaptor Ligation Mix to each sample, mixing, and incubating in the thermal cycler for 15 minutes. SPRIselect (0.8X) was used for post-ligation cleanup. Sample Index PCR, followed by a final SPRIselect (0.8X) cleanup step, completed the library preparation process.

#### Single-nuclei sequencing

Single-nuclei libraries were pooled and sequenced on the Illumina NovaSeq S2 v1.5 platform. Sequencing was paired-end using single indexing and a minimum of 20,000 read pairs per cell.

#### Bioinformatic analysis

The R package Seurat (v4.0.2) was used to generate a single-cell matrix as described previously^63^. Cycling cells were included in this analysis. Quality control (QC) filtering retained cells with feature counts >400, percentage of mitochondrial genes <1%, and percentage of ribosomal genes <5% (Supplementary Figure 5). SoupX (v1.6.2) was used to estimate and remove cell-free mRNA contamination. ParamSweep (v3.0) was used for parameter optimization and doubletFinder (v2.0.3) to remove doublets. The count matrix was normalized and 3000 variable genes were selected. After scaling, dimensionality reduction was performed using the first 30 principal components. Seurat packages FindClusters and RunUMAP were used to identify cell clusters and for Uniform Manifold Approximation and Projection (UMAP) visualization. The clustree R package (v0.5) was used to select a clustering resolution of 0.3 throughout. The 12wpc and 13wpc 46,XX ovaries were integrated for analysis using SCTransform (v0.3.5). Differential gene expression was performed using the FindAllMarkers function (‘min.pct=0.25, logfc.threshold=0.25’). Visualization functions within Seurat, including FeaturePlot, VlnPlot, and DotPlot, were used to visualize gene marker expression. All R packages and versions used in the bioinformatic analysis pipeline are described in Supplementary Methods 1.

### Statistical Analysis

Statistical analyses were performed using GraphPad Prism v9.1.1 (GraphPad Software) or in R (v4.2.0) and are described in the relevant sections. A *p* value of less than 0.05 was considered significant. The Benjamini-Hochberg approach was used to adjust for multiple testing with cut-off adjusted *p* values of 0.05^64^. Data are shown as individual data points or as violin plots where appropriate.

## Data availability and Supplementary data

### Data repository links

Bulk RNA sequencing data are deposited in ArrayExpress/Biostudies (accession number S-BSST693). Single-cell RNA sequencing data are deposited in ArrayExpress/Biostudies (accession number S-BSST1194)

### Supplementary Files

**Supplementary Movie 1:** Micro-CT of human fetal ovary and oviduct (Fallopian tube) at 20wpc

**Supplementary Materials:** Supplementary figures 1-5

**Supplementary Table 1:** Width, length, and weight of the early human fetal ovary

**Supplementary Table 2:** Top 250 differentially expressed genes, ovary v testis.

**Supplementary Table 3:** Top 250 differentially expressed genes, ovary v control.

**Supplementary Table 4:** Top 250 differentially expressed genes, testis v ovary.

**Supplementary Table 5:** Top 250 differentially expressed genes, testis v control.

**Supplementary Table 6:** Top 250 differentially expressed genes, ovary v testis 15/16wpc.

**Supplementary Table 7:** Top 250 differentially expressed genes, ovary v testis 11/12wpc

**Supplementary Table 8:** Top 250 differentially expressed genes, ovary v testis 9/10wpc.

**Supplementary Table 9:** Top 250 differentially expressed genes, ovary v testis CS22/23.

**Supplementary Table 10:** Differentially expressed genes, ovary 15/16wpc v ovary CS22/23.

**Supplementary Table 11:** The 288 highly ovary specific genes.

**Supplementary Table 12:** Comparison of differentially expressed genes (DEGs) between the developing ovary and testis

**Supplementary Table 13:** Highly ovary-specific non-coding RNA genes

**Supplementary Table 14:** Gene markers used for annotation of cell clusters

**Supplementary Table 15:** MAGE cancer-testis genes in the developing ovary.

**Supplementary Table 16:** Differentially expressed transcription factors in the fetal gonad.

**Supplementary Table 17:** Differentially expressed nuclear receptor genes, ovary v control and ovary v testis.

**Supplementary Table 18:** Genes on the “PanelApp” POI panel designed by the 100,000 Genome Study, version 1.67, August 2023.

### Author contributions

Author contributions were as follows: Study conceptualization: SMcG-B, IdV, JCA; Methodology: SMcG-B, IdV, TX, ICS, TB; Investigation: SMcG-B, IdV, TX, ICS, JPS, FB, BC, NM, DL, PN, TB, NS, JCA; Formal analysis: SMcG-B, IdV, ICS; Data curation: SMcG-B, IdV, ICS; Resources: OA, NS; Project administration: JCA; Supervision: GSC, MTD, OJA, NS, JCA; Validation: SMcG-B, IdV, TX, TB; Visualization: SMcG-B, IdV, JCA; Writing – original draft: SMcG-B, JCA; Writing – review & editing: All authors; Funding acquisition: SMcG-B, OA, JCA.

## Acknowledgements

This research was funded in whole, or in part, by the Wellcome Trust Grants 216362/Z/19/Z to SMcG-B and 209328/Z/17/Z to JCA. For the purpose of Open Access, the author has applied a CC-BY public copyright license to any Author Accepted Manuscript version arising from this submission. ICS received funding from the National Institute of Health Research (NIHR) (ICA-CDRF-2017-03-53 and NIHR302390). Human fetal material was provided by the Joint MRC/Wellcome Trust (Grant MR/R006237/1) Human Developmental Biology Resource (http://www.hdbr.org). Research at UCL Great Ormond Street Institute of Child Health is supported by the National Institute for Health Research, Great Ormond Street Hospital Biomedical Research Centre (grant IS-BRC-1215-20012). The views expressed are those of the authors and not necessarily those of the National Health Service, National Institute for Health Research, or Department of Health. The funders had no role in study design, data collection and analysis, decision to publish, or preparation of the manuscript. We thank other members of UCL Genomics and the Human Developmental Biology Resource for their additional contributions to this work. This work forms part of a PhD Thesis submitted to University College London (SMcG-B).

