## Supplemental Materials for "Mapping the anatomical and transcriptional landscape of early human fetal ovary development"

### Supplementary Methods 1:

R Packages used in scRNAseq analysis

| | | | |
|---|---|---|---|
| tidyr_1.2.1 | fansi_1.0.3 | listenv_0.9.0 | renv_0.16.0 |
| abind_1.4-5 | farver_2.1.1 | lmtest_0.9-40 | reshape2_1.4.4 |
| AnnotationDbi_1.56.2 | fastmap_1.1.0 | locfit_1.5-9.7 | reticulate_1.26 |
| AnnotationHub_3.2.2 | filelock_1.0.2 | magrittr_2.0.3 | rhdf5_2.38.1 |
| assertthat_0.2.1 | fitdistrplus_1.1-8 | MASS_7.3-58.1 | rhdf5filters_1.6.0 |
| beachmat_2.10.0 | future_1.30.0 | Matrix_1.5-3 | Rhdf5lib_1.16.0 |
| Biobase_2.54.0 | future.apply_1.10.0 | MatrixGenerics_1.6.0 | rlang_1.0.6 |
| BiocFileCache_2.2.1 | generics_0.1.3 | matrixStats_0.63.0 | ROCR_1.0-11 |
| BiocGenerics_0.40.0 | GenomeInfoDb_1.30.1 | memoise_2.0.1 | RSQLite_2.2.20 |
| BiocManager_1.30.19 | GenomeInfoDbData_1.2.7 | mime_0.12 | rstudioapi_0.14 |
| BiocParallel_1.28.3 | GenomicRanges_1.46.1 | miniUI_0.1.1.1 | Rtsne_0.16 |
| BiocVersion_3.14.0 | ggforce_0.4.1 | munsell_0.5.0 | S4Vectors_0.32.4 |
| Biostrings_2.62.0 | ggplot2_3.4.0 | nlme_3.1-161 | scales_1.2.1 |
| bit_4.0.5 | ggraph_2.1.0 | parallel_4.1.0 | scattermore_0.8 |
| bit64_4.0.5 | ggrepel_0.9.2 | parallelly_1.33.0 | sctransform_0.3.5 |
| bitops_1.0-7 | ggridges_0.5.4 | patchwork_1.1.2 | scuttle_1.4.0 |
| blob_1.2.3 | globals_0.16.2 | pbapply_1.6-0 | Seurat_4.3.0 |
| cachem_1.0.6 | glue_1.6.2 | pillar_1.8.1 | shiny_1.7.4 |
| celldex_1.4.0 | goftest_1.2-3 | pkgconfig_2.0.3 | SingleCellExperiment_1.16.0 |
| cli_3.5.0 | graphlayouts_0.8.4 | plotly_4.10.1 | sparseMatrixStats_1.6.0 |
| cluster_2.1.4 | grid_4.1.0 | plyr_1.8.8 | spatstat.data_3.0-0 |
| clustree_0.5.0 | gridExtra_2.3 | png_0.1-8 | spatstat.explore_3.0-5 |
| codetools_0.2-18 | gtable_0.3.1 | polyclip_1.10-4 | spatstat.geom_3.0-3 |
| colorspace_2.0-3 | HDF5Array_1.22.1 | progressr_0.12.0 | spatstat.random_3.0-1 |
| compiler_4.1.0 | htmltools_0.5.4 | promises_1.2.0.1 | spatstat.sparse_3.0-0 |
| cowplot_1.1.1 | htmlwidgets_1.6.0 | purrr_1.0.0 | spatstat.utils_3.0-1 |
| crayon_1.5.2 | httpuv_1.6.7 | R.methodsS3_1.8.2 | splines_4.1.0 |
| curl_4.3.3 | httr_1.4.4 | R.oo_1.25.0 | stringr_1.5.0 |
| data.table_1.14.6 | ica_1.0-3 | R.utils_2.12.2 | SummarizedExperiment_1.24.0 |
| DBI_1.1.3 | igraph_1.3.5 | R6_2.5.1 | survival_3.4-0 |
| dbplyr_2.2.1 | interactiveDisplayBase_1.32.0 | RANN_2.6.1 | tensor_1.5 |
| DelayedArray_0.20.0 | IRanges_2.28.0 | rappdirs_0.3.3 | tibble_3.1.8 |
| DelayedMatrixStats_1.16.0 | irlba_2.3.5.1 | RColorBrewer_1.1-3 | tidygraph_1.2.2 |
| deldir_1.0-6 | jsonlite_1.8.4 | R.oo_1.25.0 | tidyselect_1.2.0 |
| digest_0.6.31 | KEGGREST_1.34.0 | R.utils_2.12.2 | tools_4.1.0 |
| DoubletFinder_2.0.3 | KernSmooth_2.23-20 | R6_2.5.1 | tweenr_2.0.2 |
| dplyr_1.0.10 | labeling_0.4.2 | RANN_2.6.1 | utf8_1.2.2 |
| dqrng_0.3.0 | later_1.3.0 | rappdirs_0.3.3 | uwot_0.1.14 |
| DropletUtils_1.14.2 | lattice_0.20-45 | RColorBrewer_1.1-3 | vctrs_0.5.1 |
| edgeR_3.36.0 | lazyeval_0.2.2 | Rcpp_1.0.9 | viridis_0.6.2 |
| ellipsis_0.3.2 | leiden_0.4.3 | RcppAnnoy_0.0.20 | SoupX_1.6.2 |
| euratObject_4.1.3 | lifecycle_1.0.3 | RCurl_1.98-1.9 | ParamSweep_3.0 |
| ExperimentHub_2.2.1 | limma_3.50.3 | remotes_2.4.2 | SCTransform_0.3.5 |

**Supplementary Methods 2:**

Components for the buffers used for single-nuclei suspension:

*Salty Ez10 Lysis Buffer*

10 mM Tris-HCl pH 7.5 1M

146 mM NaCl 5M

1 mM CaCl2 1M

21 mM MgCl2 1M

0.03% Tween-20 (Sigma-Aldrich)

0.01% BSA (Miltenyi Biotec)

10% Ez Lysis Buffer (Sigma-Aldrich)

0.2-1 U/uL Protector RNAse Inhibitor (Roche)

1 mM DTT (ThermoFisher Scientific)

*Wash and Resuspension Buffer 2 (WRB2)*

10 mM Tris-HCl pH 7.5

10 mM NaCl

3 mM MgCl2

1mM DTT (ThermoFisher Scientific)

1% BSA (Miltenyi Biotec)

0.2-1 U/uL Protector RNAse Inhibitor (Roche)

**Supplementary Figure 1. Bulk RNA sequencing experimental design**

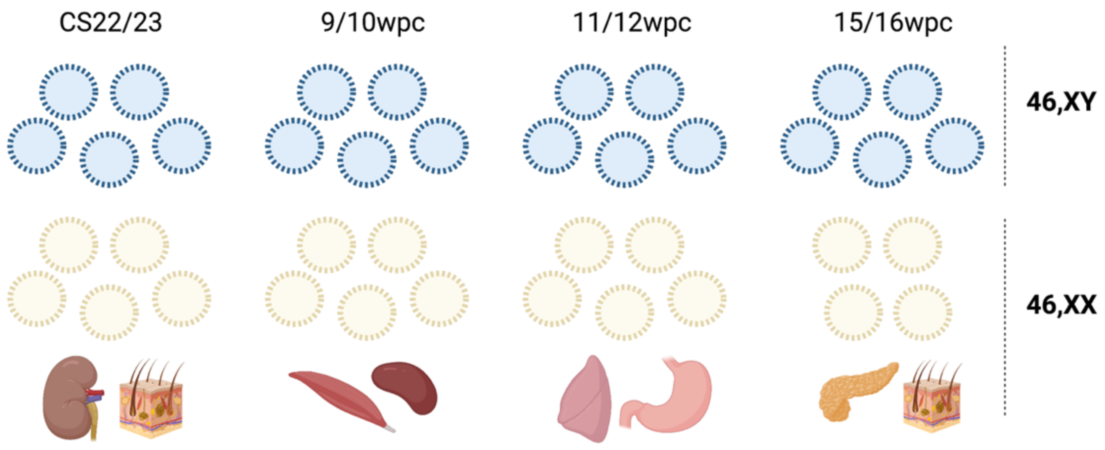


Testis (blue), ovary (yellow), and 46,XX control samples (kidney, skin, muscle, spleen, lung, stomach, pancreas) were collected for sequencing from each of four key developmental stages (CS22/23; 9/10wpc; 11/12wpc; 15/16wpc).

**Supplementary Figure 2. Cluster analysis of experimental samples.**

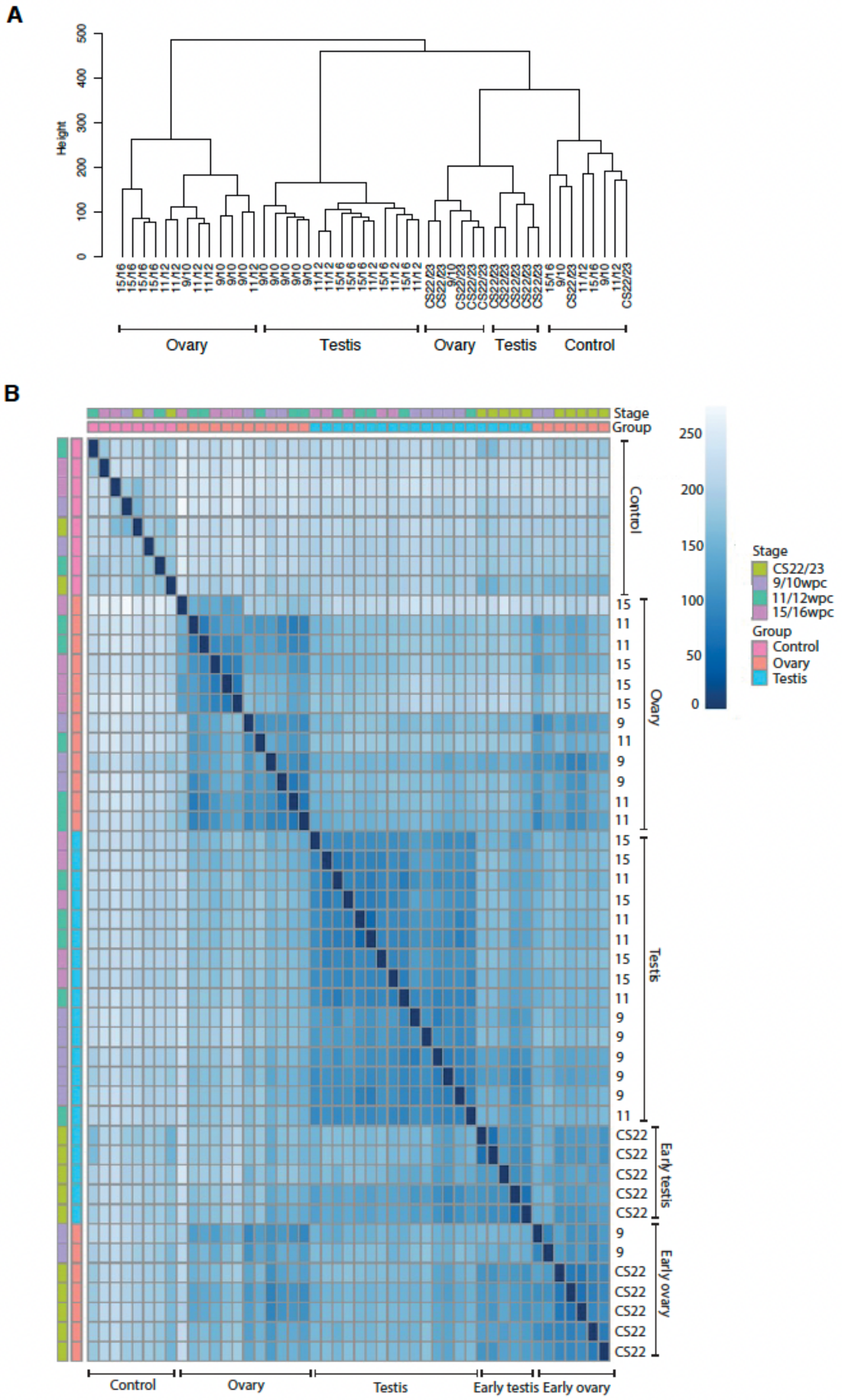

**A)** Cluster dendrogram of all 47 samples used in the study demonstrating clustering by developmental stage and tissue type. The hierarchical agglomeration clustering method (Ward's method, or ward.D2) was used for analysis**. B)** Correlation heatmap of gene expression across all 47 samples. Darker intensity and scores closer to 0 indicate higher correlation between samples.

**Supplementary Figure 3. Non-coding transcripts in the developing fetal gonad.**

**A**

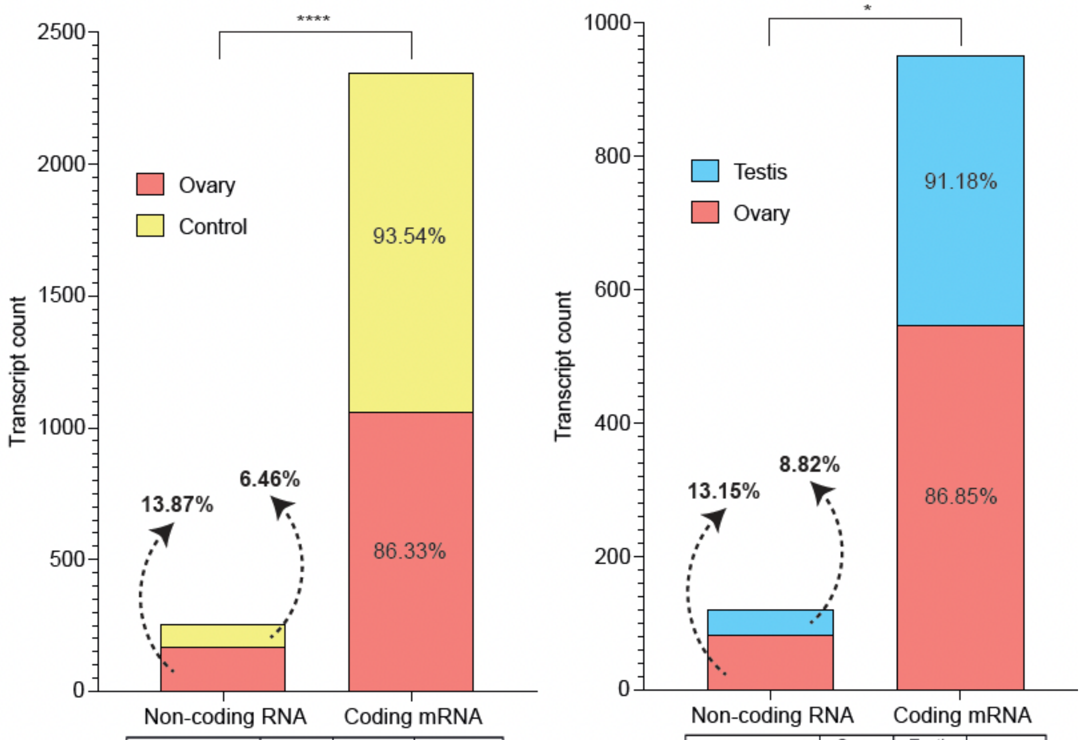


| | *Ovary* | *Control* | *p* |
|---|---|---|---|
| **Non-coding** | 168 | 89 | **<0.0001** |
| **Coding** | 1061 | 1289 | |
| **Total** | 1229 | 1378 | |

| | *Ovary* | *Testis* | *p* |
|---|---|---|---|
| **Non-coding** | 83 | 39 | **0.0314** |
| **Coding** | 548 | 403 | |
| **Total** | 631 | 442 | |

**B**

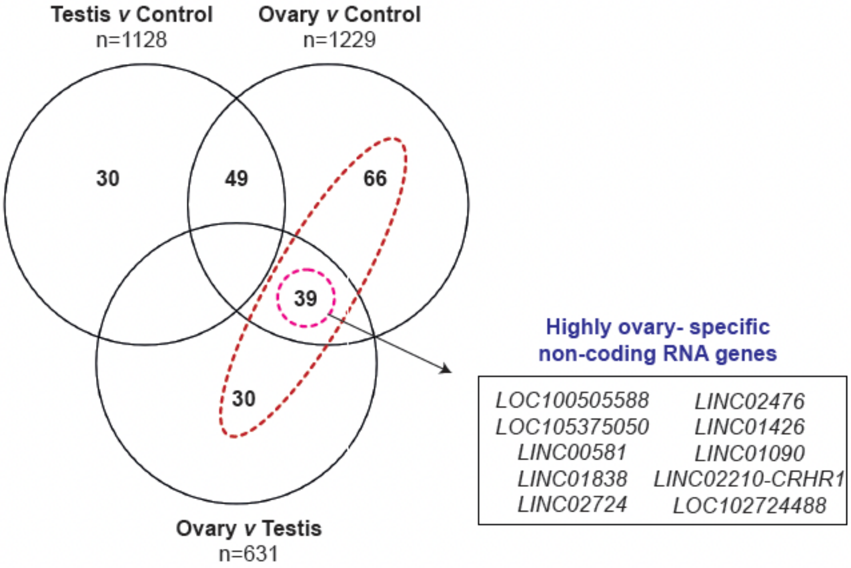

**A)** Total numbers of differentially expressed non-coding and coding transcripts between the ovary and control (left panel) and the ovary and testis (right panel) were examined globally ($log_2FC>2$; p.adj<0.05). Percentages of non-coding and coding transcripts compared to the total number of differentially expressed genes in the two analyses are indicated on both graphs. Differences in proportions of non-coding transcripts in the ovary compared to control or testis were examined using a Fisher's exact test (*$p<0.05$; ****$p<0.0001$). **B)** Absolute numbers of differentially expressed non-coding transcripts ($log_2FC>2$; p.adj<0.05) in the ovary compared to control; the testis compared to control; and the ovary compared to testis are shown. Ovary-specific (red) and highly ovary-specific transcripts (pink) are indicated. The top 10 differentially expressed highly ovary-specific non-coding RNA transcripts are indicated on the Venn diagram.

**Supplementary Figure 4. Pathway enrichment analysis of genes differentially expressed in the ovary compared to control and to testis.**

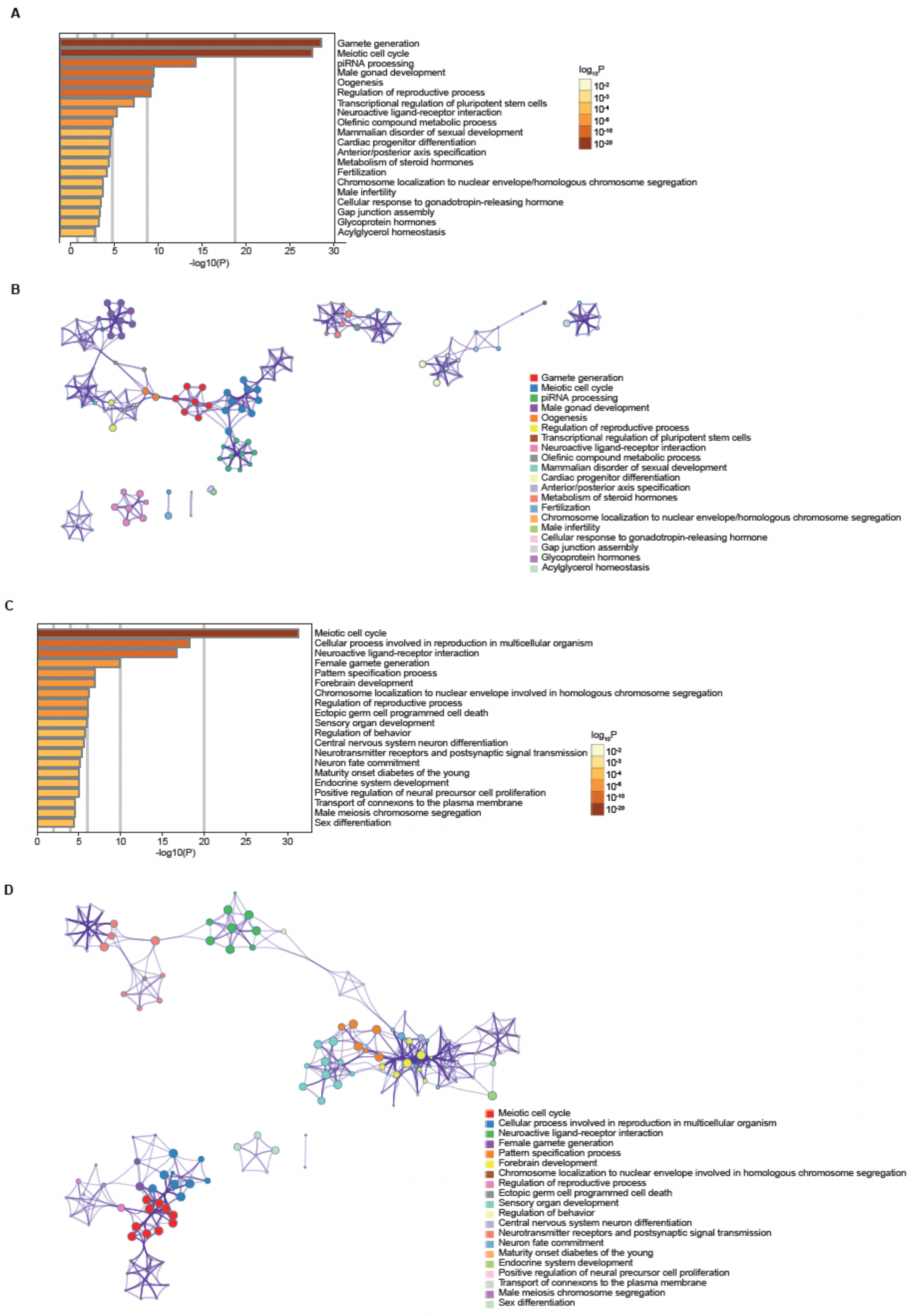

**A)** Bar graph of enriched gene ontology terms within the ovary *v* control analysis with a $\log_2FC>2$, p.adj<0.05; n=1229 total genes). Bars are coloured by $\log_{10}p$ value with darker coloured bars indicating greater significance. **B)** Network analysis and visualisation of enriched gene ontology terms in the in the ovary *v* control analysis where each node represents an enriched term coloured by cluster ID (Metascape). Nodes with the same cluster ID cluster together. Terms with >0.3 similarity are connected by edges. **C)** Bar graph of enriched gene ontology terms within the ovary *v* testis analysis with a $\log_2FC>2$, p.adj<0.05; n=631 total genes). **D)** Network analysis and visualisation of enriched gene ontology terms in the ovary *v* testis analysis.

**Supplementary Figure 5: snRNAseq RNA sequencing quality control**

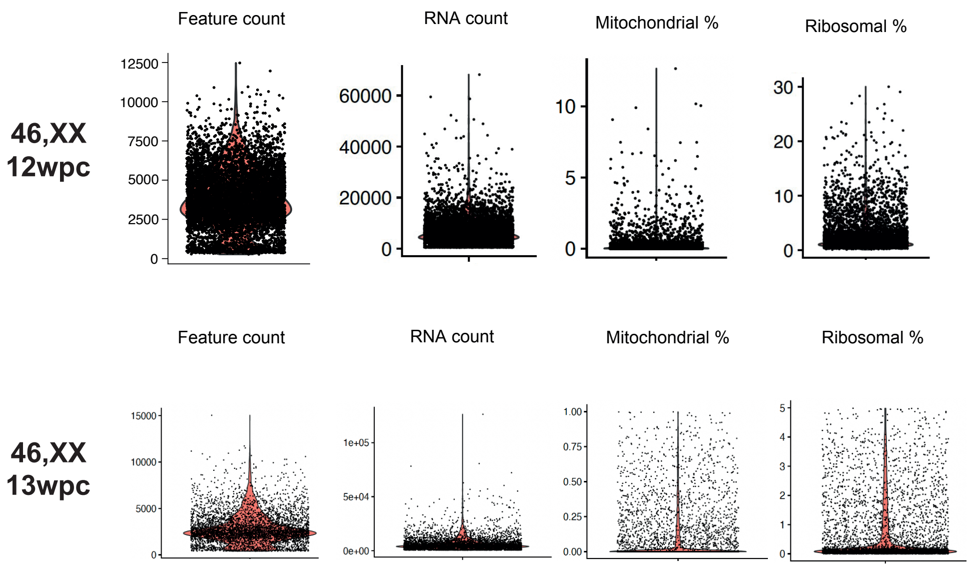


| | *13wpc 46,XX* | *12wpc 46,XX* |
|---|---|---|
| Raw cells | 5121 | 7033 |
| <1% mitochondrial counts | 4884 | 6745 |
| <5% ribosomal counts | 4732 | 6180 |
| Unique feature counts >400 | 4705 | 6179 |
| Doublets removed | 4489 | 5802 |
| **Final cell count** | **4489** | **5802** |

**Upper panel:** For each of the four samples included in the snRNAseq analysis, the feature count, RNA count, percent mitochondrial genes and percent ribosomal genes are shown.

**Lower panel:** The starting number of raw cells is shown for each ovary sample. Cells were then removed from the analysis if they had >1% mitochondrial counts, >5% ribosomal counts, unique feature counts <400, or were identified as doublets. The final cell count for analysis is shown in bold.

**Supplementary Figure 6: Single-nuclei expression of *THRA* and *THRB***

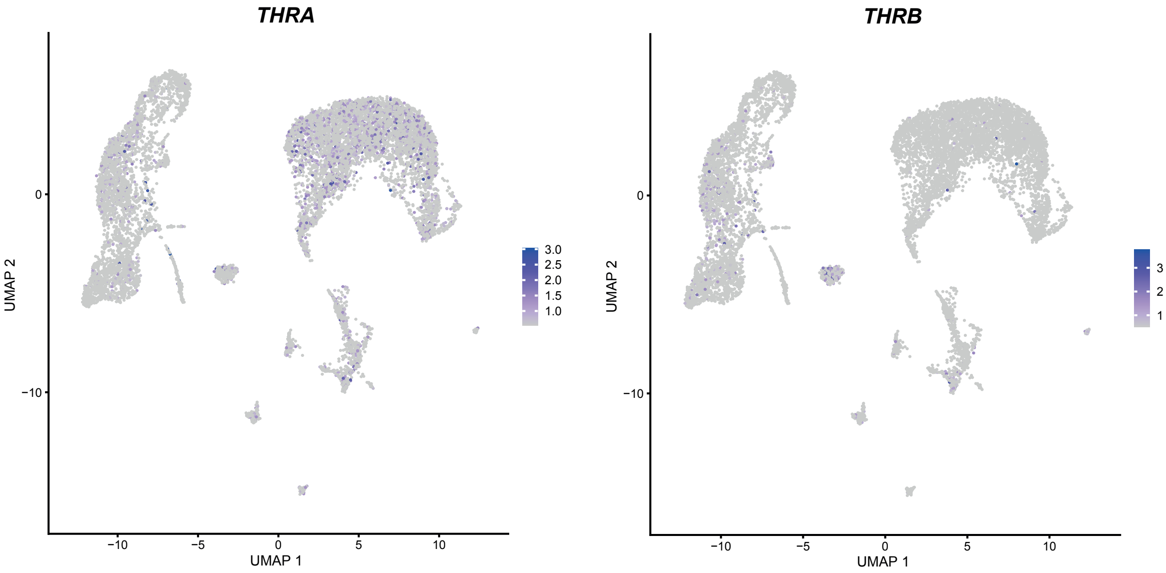


UMAP representation of the thyroid hormone receptors *THRA* and *THRB* in the snRNAseq data demonstrating a particular localisation of *THRA* to pre-granulosa cell populations.
